# Immune repertoire profiling reveals extensive clonal expansion of T and B cells in CXCL9-rich niches in sinonasal cancer

**DOI:** 10.64898/2026.06.04.730068

**Authors:** Javiera A. Álvarez, Eva Gracia Villacampa, Daniel Labuz, Mariia Guryleva, Egon Urgard, Åsa Kågedal, Oscar Solmell, Anders Näsman, Javier Escudero Morlanes, Junjie Ma, Niels Ødum, Adnane Achour, Susanna Kumlien Georén, Benedict Chambers, Benjamin Murrell, Lars Olaf Cardell, Camilla Engblom, Kim Thrane, Jonathan M. Coquet

## Abstract

Sinonasal cancer (SNC) is a rare and aggressive head and neck cancer with limited therapeutic options and an incompletely defined immune microenvironment. Although histopathology remains central to clinical tumor evaluation, immune profiling typically requires tissue dissociation and therefore loses the spatial relationships that shape antitumor immunity. Here, we combined single-cell RNA-sequencing approaches, spatial transcriptomics and Spatial V(D)J, a technique we recently pioneered, to analyse the cellular, spatial, and clonal architecture of SNC. We found that SNC tumors were marked by a robust T cell infiltration, including substantial proportions of classical tissue resident memory (TRM) and exhausted (TExh) CD8^+^ T cells. This inflamed phenotype was accompanied by an immense infiltration of suppressive regulatory T (Treg) cells, which differentiated towards a Tbet^+^CXCR3^+^ phenotype. Spatially, expanded Treg, CD8⁺ T and B cell clones concentrated within specialized peritumoral immune niches enriched for cancer-associated fibroblasts, CXCL9⁺ tumor-associated macrophages and LAMP3⁺IDO1⁺ dendritic cells. T cell clones occupying these niches were clonally related to those infiltrating epithelial tumor regions, linking lymphoid hubs to the broader tumor immune response. Together, these data identify SNC as an inflamed but highly immunoregulatory tumor type and reveal a spatially organized clonal architecture in which suppressive and cytotoxic lymphocyte states coexist within CXCL9⁺ myeloid niches.

## Introduction

Solid cancers are spatially organized ecosystems composed of malignant and non-malignant cells, embedded within extensive extracellular matrix and vascular networks. Successful cancers reprogram the extracellular environment, known as the tumor microenvironment (TME), to promote their growth (Hanahan and Weinberg 2011), and evade immune-mediated elimination.

While the TME comprises a range of cell types including the tumor itself, fibroblasts, macrophages, innate lymphocytes, B and NK cells, the presence of T cells is a key determinant of tumor progression or eradication. T cell-driven tumor immunity is closely linked to tumor mutational burden and likely depends on the presence of tumor ‘neoantigens’ that are recognized by T cells through the T cell antigen receptor (TCR) (Schumacher and Schreiber 2015). The balance between cytotoxic CD8⁺ T lymphocytes with tumor-killing capacity, and suppressive FOXP3^+^ regulatory CD4^+^ T cells (Tregs), has long been proposed as a major factor influencing tumor growth and patient outcome (Saleh and Elkord 2020; Galon et al. 2006; Fridman et al. 2012; Sato et al. 2005). However, this dichotomy is too simplistic to fully explain the function of T cells in solid cancers.

CD8^+^ T cells encompass several transcriptionally and functionally-distinct states, including stem-like, cytotoxic, tissue-resident memory (TRM) and exhausted (TExh) cells. These states share overlapping transcriptional programs in non-lymphoid tissues and in solid tumors, complicating their distinction (Wijesinghe et al. 2025; Burn et al. 2026; Masopust et al. 2025). Recent studies that refined the transcriptional signatures of TExh and TRM cells in solid tumors found that TRM cells were associated with better patient prognosis, while TExh cells could be reinvigorated by PD-1 blockade to improve patient outcomes (Savas et al. 2018; Gavil et al. 2024; Wu et al. 2025; Burn et al. 2026; Park et al. 2026). Similarly, CD4^+^ T cells can be found in distinct stem-like, cytotoxic and Treg cell states. In solid cancers, Treg cells have been shown to acquire cancer-specific adaptations including expression of CCR8 and IL1R1, which are associated with their persistence and suppressive function (Villarreal et al. 2018; Kidani et al. 2022; Mair et al. 2022), while stem-like CD4^+^ T cells can give rise to Th1 effectors and enhance CD8^+^ T cell-mediated tumor immunity (Cardenas et al. 2024).

Despite an increasingly granular understanding of the function of specific T cell states, T cell clonal evolution and the spatial relationship between T cell states remains largely unclear, due to technical limitations in tracking cell states and clones directly in tumor tissue. Preserving the spatial organisation of the tumor may identify clones positioned to engage malignant cells, reveal spatial associations between CD4^+^ and CD8^+^ T cell clonotypes, and define tumor niches that support distinct T cell clones or states.

In this study, we profile the immune and spatial composition of sinonasal squamous cell carcinoma (SNC), a rare head and neck cancer characterized by aggressive local growth, limited therapeutic options, and poor prognosis. SNC is typically HPV-negative (Tendron et al. 2022) and infrequently displays lymph node involvement at the time of diagnosis. Despite its clinical severity, SNC remains poorly understood at the molecular and immunological levels (Esposito et al. 2022). Combining single-cell RNA-Sequencing (scRNA-seq), spatial transcriptomics, our Spatial V(D)J workflow (Engblom et al. 2023) and a newly developed PhyloTrajectory tool, we investigate the architecture of SNC at the clonal level. Our findings reveal an extensive evolution of the adaptive lymphocyte compartment in SNC and an intimate spatial relationship between hundreds of T and B cell clones simultaneously that may have implications for solid cancers in general.

## Results

### SNC tumors exhibit heterogeneous yet spatially organized histopathological and immune architectures

To interrogate SNC in detail, we analyzed surgically resected tumors from 9 patients using a combination of scRNA-seq, spatial transcriptomics, immune repertoire profiling, flow cytometry and fluorescence microscopy (Fig 1A, Supplementary Table 1). Three tumors were spatially profiled using the 10x Genomics Visium platform, yielding 8 tissue sections and 13,810 spatial spots across SNC2, SNC3 and SNC4. Quality-control metrics showed consistent RNA capture across sections, supporting robust spatial profiling across replicate sections (Supplementary Fig. 1A).

**Figure 1.**
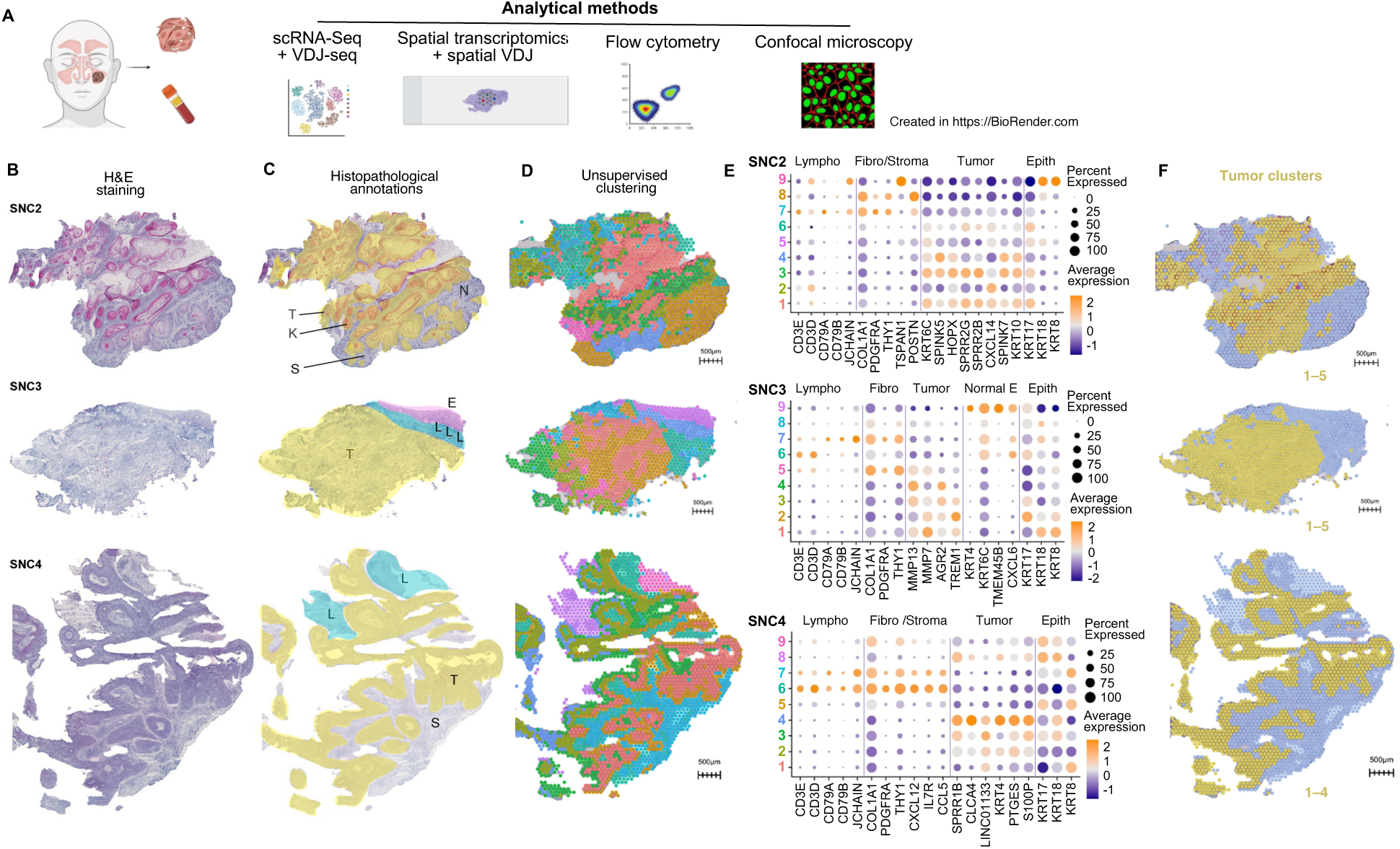
Spatial transcriptomics reveals organized tumor, stromal and immune compartments in sinonasal cancer. (**A**) Schematic overview of the multi-modal experimental approaches used in this study. Tumor tissue and peripheral blood from SNC, HNSCC patients and healthy donors were profiled using scRNA-seq with V(D)J sequencing, spatial transcriptomics with spatial V(D)J profiling, flow cytometry, and confocal imaging. Supplementary table 1 outlines which analyses were undertaken on each tumor. (**B**) H&E staining of representative sections from three SNC tumors (SNC2-4). (**C**) Pathologist-derived tissue annotations highlighting major histological compartments including tumor (T), stroma (S), keratin pearls (K), necrosis (N), lymphoid areas (L), and epithelium (E). (**D**) Spatial transcriptomic unsupervised clustering map of SNC2-4. (**E**) DotPlots display canonical markers for lymphocytes, fibroblasts/stroma, and epithelial cells. Tumor markers correspond to the differentially expressed genes enriched in histologically defined tumor regions (yellow color in panel C). Cluster numbers are color-coded to match the corresponding clusters shown in panel D. (**F**) Spatial maps of merged tumor clusters in each tumor. All scale bars, 500 µm.

Histopathological examination revealed distinct tumor architectures across patients, including keratinizing squamous cell carcinoma in SNC2, poorly differentiated non-keratinizing tumor architecture in SNC3, and papillary non-keratinizing tumor architecture in SNC4 (Fig. 1B-C; Supplementary Note 1). Consistent with these morphological differences, spatial transcriptomic profiles showed pronounced inter-patient heterogeneity, with UMAP visualization separating sections primarily by patient identity (Supplementary Fig. 1B). We therefore performed unsupervised clustering independently for each patient, while replicate sections from the same tumor showed consistent transcriptional organization (Supplementary Fig. 1C-D).

Spatial clustering resolved patient-specific tumor, stromal and immune-enriched regions (Fig. 1D-E, Supplementary Fig. 1C,E). Across tumors, epithelial tumor regions were marked by keratin-associated and tumor-associated genes, whereas lymphocyte-rich stromal regions expressed T-cell, B-cell and plasma-cell markers, including CD3 genes, *CD79A* and *JCHAIN*. These immune-enriched regions frequently coincided with fibroblast and extracellular-matrix-associated signals, including *POSTN*, *THY1* and *COL1A1*, indicating spatial coupling between lymphoid infiltration and fibroblast-rich stromal remodeling.

Thus, despite marked inter-patient heterogeneity, all profiled SNC tumors displayed a spatially organized tumor–stroma–immune architecture, with B- and T-cell-rich stromal niches positioned near tumor regions and fibroblast-activated zones. To compare transcriptomically defined tumor regions with histopathological annotations, clusters with predominant epithelial tumor features were merged into broader tumor groups. These broadly matched the histophathologically annotated tumor regions, while retaining some section-specific differences (Fig. 1F).

### SNC are composed of conventional αβ T cells, γδ T cells and NK cells

To complement the spatial and histological analyses, we performed scRNA-seq on seven tumors to characterize the cellular composition of SNC. Cell sorting was biased toward T cells, given their relevance to antitumor immunity, but also captured other CD45^+^ and CD45^-^ cells (Supplementary Fig. 2A-D). After quality control, normalization, and integration using Seurat (Stuart et al. 2019), a total of ∼21,000 single-cell transcriptomes were retained. Unsupervised clustering revealed αβ T cells, B cells, plasma cells, epithelial cells, mast cells, myeloid cells, neutrophils, natural killer (NK) cells and γδ T cells (Supplementary Fig. 2A-F). All tumors were classified as primary SNC, except tumor 6, later confirmed as a cutaneous squamous cell carcinoma secondarily involving the sinonasal cavity. As its exclusion did not alter clustering or marker genes (Adjusted Rand Index = 1.0; mean Jaccard for all cells = 0.88; Supplementary Fig. 3A-C), it was retained in the integrated analysis. We next generated lineage-focused UMAPs for CD8⁺ and CD4⁺ T cells using an iterative subclustering approach (Supplementary Fig. 2A-F, Methods).

### CD8^+^ T cells exist on a spectrum of states including stem-like, cytotoxic, TRM and TExh in SNC

Analysis of intratumoral CD8⁺ T cells identified nine transcriptionally distinct clusters (Fig. 2A). To characterize CD8⁺ T cell states, we applied manually curated gene signatures representing stem-like/memory, cytotoxic, TRM, TExh and dividing phenotypes, and calculated enrichment scores using Seurat’s *AddModuleScore* (Fig. 2C-D, Supplementary Table 2). Across tumors, features of stem-like, cytotoxic, TRM and TExh cells were consistently detected (Fig. 2C). In general, annotations were intuitive, with clusters biased towards certain gene signatures, although the annotated TRM 2 cluster also contained some features of exhaustion (Fig. 2C).

**Figure 2.**
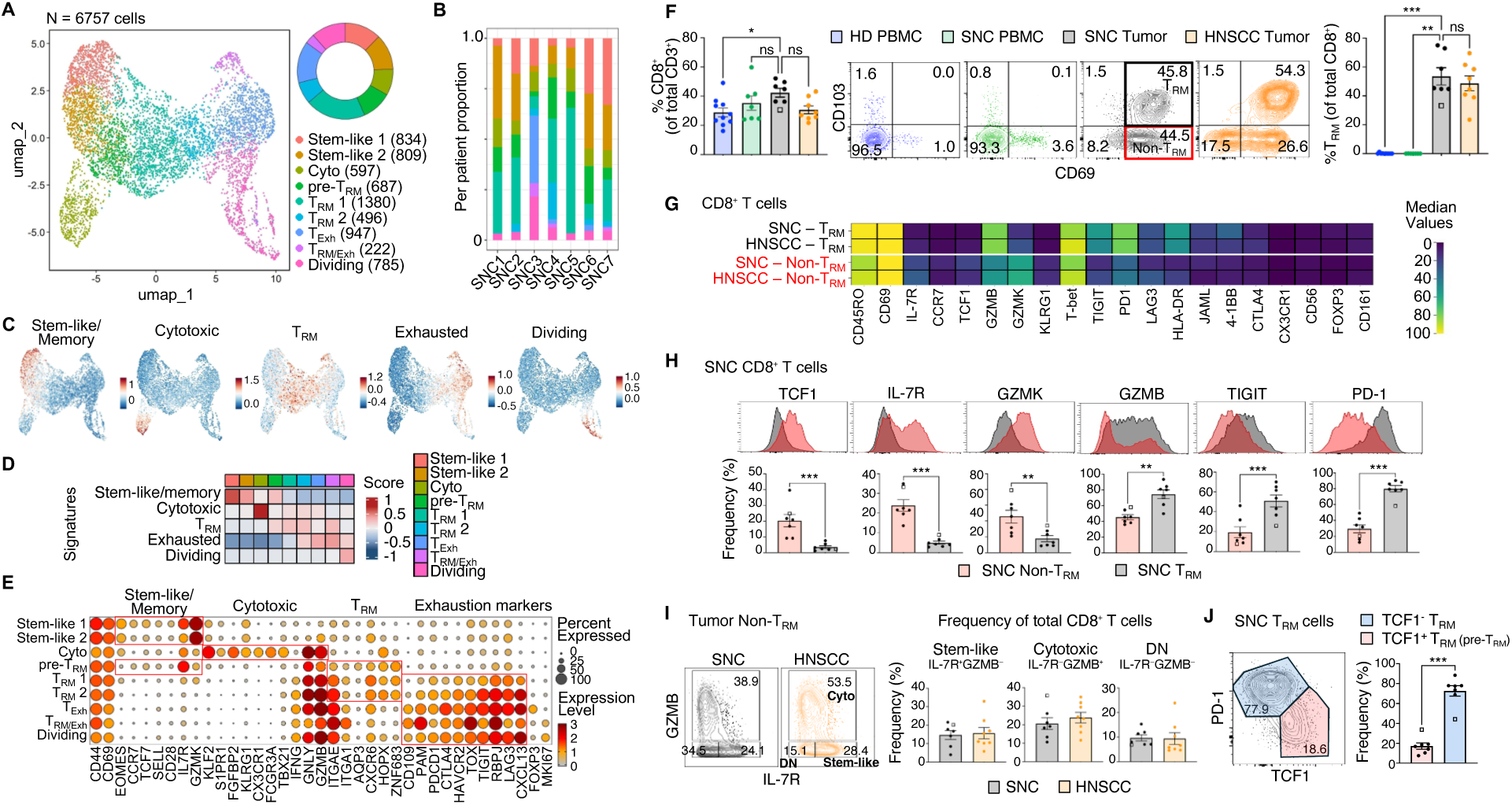
CD8⁺ T cells in SNC comprise stem-like, cytotoxic, TRM and TExh subsets. (**A**) UMAP visualization of 6,757 CD8⁺ T cells integrated across SNC tumors. The donut chart indicates the relative abundance of each cluster in the combined dataset. (**B**) Distribution of CD8⁺ T cell clusters across individual SNC tumors. (**C**) UMAP projections of CD8⁺ T cells showing module scores for manually curated gene signatures corresponding to stem-like/memory, cytotoxic, TRM, TExh, and dividing cells (scaled expression). (**D**) Heatmap of average module scores for manually curated CD8⁺ T cell gene programs across clusters. (**E**) DotPlot showing the average expression and percentage of cells expressing canonical stem-like/memory, cytotoxic, TRM, and exhaustion-associated genes across CD8⁺ T cell clusters. (**F**) Flow cytometric analysis of CD8⁺ T cells in healthy donor blood (HD PBMC), SNC blood (SNC PBMC), and tumor issue (SNC and HNSCC). Left, frequency of CD8⁺ T cells. Middle, representative FACS plots of CD69 vs. CD103 expression in CD8⁺ T cells. Right, frequency of TRM (CD69⁺CD103⁺) cells. Statistical significance was assessed using a Kruskal-Wallis test followed by multiple-comparison correction. (**G**) Heatmap of protein expression levels on gated TRM or non-TRM (CD103⁻CD69⁺) cells, median percentage values shown (SNC n = 7, HNSCC n = 8). (**H**) Representative flow cytometry histograms (top) and bar plots (bottom) of the indicated markers in tumor-infiltrating CD8⁺ TRM and non-TRM cells. Statistical significance was assessed using a paired t-test. (**I**) Representative flow cytometry plots (left) of IL-7R vs. GZMB in tumor-infiltrating non-TRM CD8⁺ T cells. Bar plots indicate the frequency of the indicated population relative to total CD8⁺ T cells, whereas flow cytometry plots show frequencies within the non-TRM gate. (**J**) Representative flow cytometry plot (left) of TCF1 vs. PD-1 expression in TRM CD8⁺ T cells from SNC. Bar graphs (right) represent mean ± SEM. Individual points indicate biological replicates; the square indicates SNC6, a cutaneous carcinoma with nasal extension. Statistical significance was assessed using a paired t-test.

Two clusters, termed Stem-like 1 and Stem-like 2, expressed *GZMK*, *CCR7*, *TCF7* and *IL7R*, along with activation markers like *CD69* and *CD44*, consistent with an activated, non-naïve stem-like CD8⁺ T-cell phenotype (Fig. 2E). This phenotype has been associated with sustained long-term antitumor immunity, gives rise to more differentiated states under chronic antigen stimulation, and is critical for responsiveness to immune checkpoint blockade (Siddiqui et al. 2019; Im et al. 2016; Kaech and Cui 2012; Miller et al. 2019).

A distinct CD8⁺ T cell population was highly enriched for cytotoxic genes, including *CX3CR1*, *KLRG1*, *TBX21*, *GZMB*/*GNLY* and *S1PR1 (Zander et al. 2019)* (Fig. 2E). This profile resembles CX3CR1⁺KLRG1⁺ TEMRA-like cytotoxic populations described in colorectal cancer, which express homing receptors such as S1PR1/S1PR5 and may be associated with tissue egress and circulating memory cells (Zhang et al. 2018). SNC cytotoxic CD8⁺ T cells also expressed *KLF2*, a transcriptional regulator that maintains effector identity, limits entry into exhaustion and enables T-bet-driven effector cell differentiation (Fagerberg et al. 2025). Thus, SNC contain cytotoxic CD8⁺ T with putative anti-tumor potential, but their resident status in the tissue is unclear.

### A continuum of TRM cell states is observed in SNC

CD8⁺ TRM have been implicated in promoting antitumor immunity, with multiple studies linking CD103⁺ TRM-like populations to improved clinical outcomes (Savas et al. 2018; Okła et al. 2021; Damei et al. 2023). In parallel, sustained antigen stimulation can drive CD8⁺ T-cell exhaustion, and overlapping marker expression between TRM and TExh states complicates their discrimination, which motivated us to use an integrative signature-based analysis to resolve these phenotypes (Burn et al. 2026; Masopust et al. 2025; Siddiqui et al. 2019).

We identified five CD8⁺ T cell clusters which shared a gene expression program and seemed to form a continuum from tissue residency to terminal exhaustion (Fig. 2D-E, Supplementary Fig. 2H). Three clusters, termed pre-TRM, TRM1 and TRM2, expressed canonical resident-memory markers (*ITGAE*/CD103, *CD69*, *ZNF683, HOPX*) together with effector genes (*GZMB*, *GNLY*). Notably, pre-TRM cells showed the highest IL7R expression, which progressively declined across TRM1 and TRM2 states. In parallel, GZMK and TCF7 expression were reduced across the pre-TRM, TRM1 and TRM2 states, while exhaustion-associated genes such as PDCD1, CTLA4 and TOX increased (Fig. 2E). TRM cells were present in all tumors examined (Fig. 2B), suggesting that the SNC TME supports their long-term T cell retention and adaptation.

A TExh population expressed high levels of *PDCD1*, *CTLA4*, *TOX*, *CD109* and *TIGIT,* maintained some TRM markers including *CD69* and *ITGAE*, but lost *HOPX* and *ZNF683* expression. This cluster also expressed *FOXP3* (Fig. 2E) which, although classically associated with CD4^+^ Tregs, can be transiently induced in activated CD8⁺ T cells (Lozano et al. 2022), with unclear functional significance in this context. TExh cells were strongly enriched in SNC3 (Fig. 2B), the only SNC patient currently deceased in our cohort, suggesting that extensive CD8⁺ T cell exhaustion may be associated with more advanced or aggressive tumor states, although additional cases are needed to determine its clinical relevance.

A small TRM/Exh cluster co-expressed tissue-residency and exhaustion markers, including elevated *TOX* together with *RBPJ*, consistent with tissue-resident exhausted states described in chronic settings (Milner et al. 2020; Pallett et al. 2017). Notably, this population showed lower *PDCD1*, *LAG3*, and *TIGIT* expression than TExh cells, suggesting a resident-adapted exhausted state with distinct checkpoint-receptor engagement (Fig. 2E).

Finally, a dividing cluster was characterized by strong *MKI67* expression, indicative of active cell cycling within the CD8^+^ T cell compartment.

### The phenotype and frequency of Stem-like, TRM and cytotoxic CD8⁺ T-cell subsets in SNC is mirrored in other HNSCC

To validate the transcriptionally defined CD8⁺ T cell subsets, we performed multiparametric flow cytometry on cryopreserved tumors and matched peripheral blood samples from SNC and HNSCC, alongside healthy donor blood (Fig. 2F-J). These included previously analyzed patients for whom tumor material was not limiting, as well as additional patients (SNC8-9 and HNSCC1-8, supplementary table 1).

Across tumors, TRM (CD103⁺CD69⁺) cells represented a major fraction of intratumoral CD8⁺ T cells (mean ∼50%) and cells of this phenotype were largely absent from circulation (Fig. 2F). TRM frequencies varied between individual tumors (from ∼30% to >70% of CD8⁺ T cells), with no clear differences in frequency between SNC and HNSCC.

Phenotypic profiling revealed that TRM cells expressed higher levels of inhibitory and activation-associated markers, including PD-1, TIGIT, LAG3, CTLA-4, JAML and GZMB, compared with non-TRM cells (CD69⁺CD103⁻) (Fig. 2G-H). In contrast, non-TRM CD8⁺ T cells were enriched for memory- and progenitor-associated markers such as IL-7R, CCR7, TCF1 and GZMK, consistent with a less differentiated or stem-like state. These patterns mirrored the scRNA-seq signatures and supported a model in which TRM cells represent a functional (GZMB^+^) and highly activated (PD-1^+^TIGIT^+^) T cell state.

Within the non-TRM fraction, co-staining for IL-7R and GZMB resolved three subsets: a cytotoxic population (GZMB⁺IL-7R⁻), a stem-like population (IL-7R⁺GZMB⁻), and a smaller double-negative population (IL-7R⁻GZMB⁻) (Fig. 2I). All three subsets were detected in both SNC and HNSCC tumors, indicating conservation of non-TRM CD8⁺ T-cell states across tumor types. Stem-like, cytotoxic and double-negative cells accounted for ∼15%, ∼20% and ∼10% of intratumoral CD8⁺ T cells, respectively and marker expression within these populations was quite similar between SNC and HNSCC (Fig. 2I and Supplementary Fig. 2I).

We also analyzed the TRM population in SNC for the presence of pre-TRM cells. We found that approximately 15% of TRM cells expressed TCF1 with low levels of PD-1 (Fig. 2J). These TCF1^+^ TRM were also enriched for IL-7R expression, while they expressed lower levels of T-bet, TIGIT and GZMB than their TCF1^-^ TRM counterparts (Supplementary Fig. 2J). This supports the possibility that a fraction of TRM cells are primed for TRM differentiation in the SNC TME.

Together, these data indicate that SNC contains verifiable stem-like, cytotoxic and TRM cells that align with the relative abundance of these cell populations in other HNSCCs.

### Regulatory T cells are a major component of the CD4^+^ T cell compartment in SNC

Phenotypic profiling of intratumoral CD4⁺ T cells identified eight transcriptionally distinct clusters (Fig. 3A-B). Using curated gene signatures and Seurat’s *AddModuleScore*, clusters were classified as stem-like (1-2), cytotoxic, Treg (1-4) and dividing populations (Fig. 3C-D, Supplementary Table 2). Two clusters, termed Stem-like 1 and Stem-like 2, expressed naïve/stem-memory-associated genes, including *SELL, CCR7*, *TCF7*, *ANXA1*, *TC2N* and *GIMAP7*, consistent with early differentiation and recirculatory potential (Fig. 3E, Supplementary Fig. 4A). Stem-like 2 additionally expressed *FOXP3* and *IKZF2* (HELIOS) at modest levels (Fig. 3E), suggestive of a stem-like population with regulatory T cell potential.

**Figure 3.**
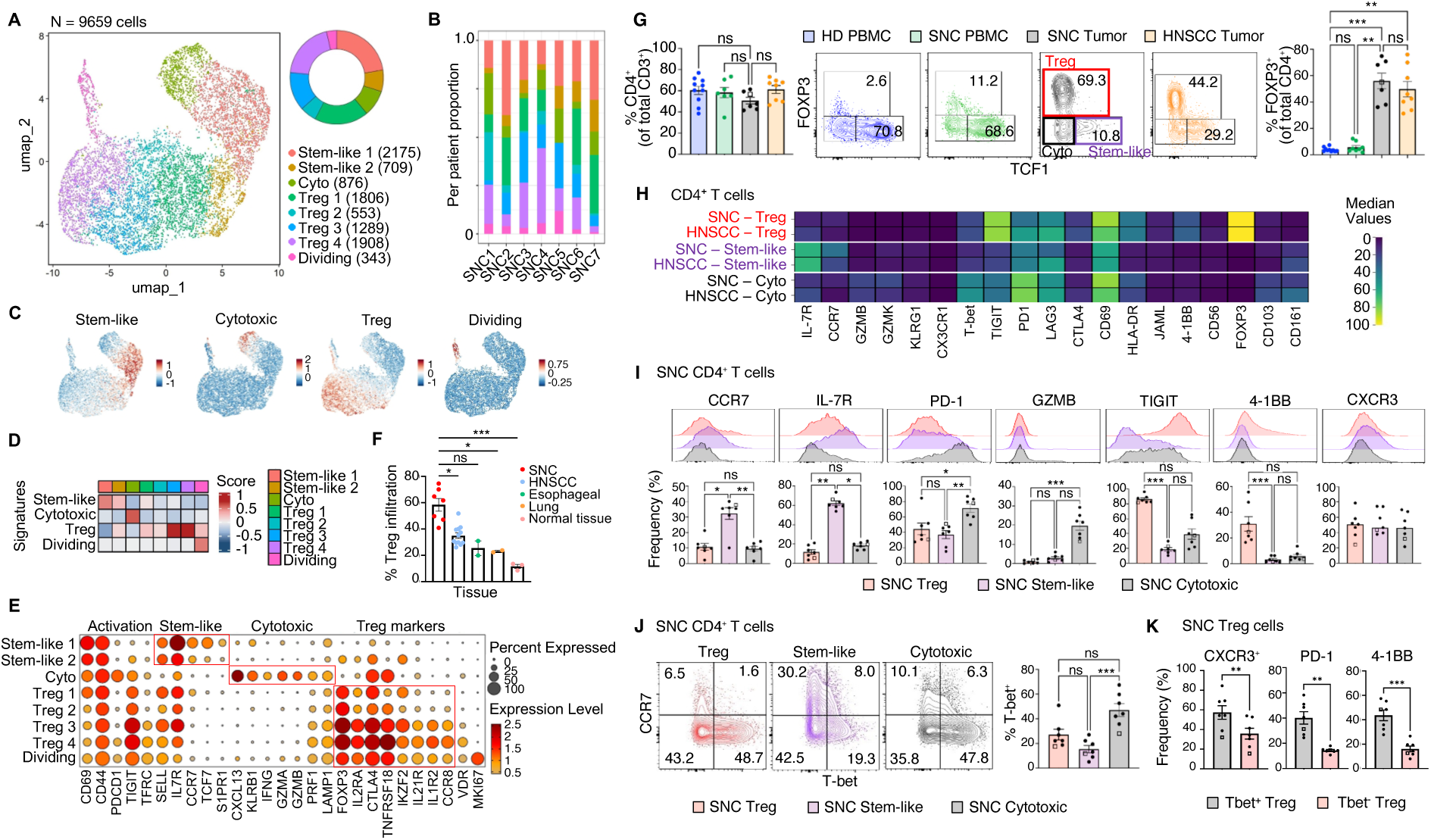
CD4⁺ T cells in SNC are primarily of a FOXP3^+^ regulatory phenotype. (**A**) UMAP visualization of 9,659 CD4⁺ T cells integrated across SNC tumors. The donut chart indicates the relative abundance of each state in the combined dataset. (**B**) Distribution of CD4⁺ T cell clusters across individual SNC tumors. (**C**) UMAP projections of CD4⁺ T cells showing module scores for manually curated gene signatures corresponding to stem-like, cytotoxic, Treg, and dividing cells. (**D**) Heatmap of average module scores for manually curated CD4⁺ T cell gene programs across clusters. (**E**) DotPlot showing the average expression and percentage of cells expressing canonical activation, stem-like, cytotoxic and Treg cell-associated genes across CD4⁺ T cell clusters. (**F**) Proportion of Treg cells among total CD4⁺ T cells across SNC, HNSCC, esophageal cancer, lung cancer, and normal tissue. Statistical significance was assessed using a Kruskal-Wallis test followed by Dunn’s multiple-comparison test. (**G**) Flow cytometric analysis of CD4⁺ T cells in healthy donor blood (HD PBMC), SNC blood (SNC PBMC), and tumor issue (SNC and HNSCC). Left, frequency of CD4⁺ T cells. Middle, representative flow cytometry plots of TCF1 vs. FOXP3 in CD4⁺ T cells, distinguishing Treg (FOXP3⁺TCF1⁻), stem-like (TCF1⁺FOXP3⁻), and double-negative populations. Right, frequency of FOXP3⁺ cells among CD4⁺ T cells. Statistical significance was assessed using a Kruskal-Wallis test followed by multiple-comparison correction. (**H**) Heatmap of protein expression levels on gated tumor-infiltrating CD4⁺ T cell subsets, median percentage values shown (SNC n = 7, HNSCC n = 8). (**I**) Representative flow cytometry histograms (top) and bar plots (bottom) of the indicated markers in tumor-infiltrating CD4⁺ Treg, stem-like, and cytotoxic populations from SNC. Statistical significance was assessed using a Friedman test followed by multiple-comparison correction. (**J**) Representative flow cytometry plots (left) of T-bet vs. CCR7 in stem-like, cytotoxic, and Treg cells. Bar plots (right) show the frequency of T-bet^+^ cells. Statistical significance was assessed using a Friedman test followed by multiple-comparison correction. (**K**) Flow cytometric analysis of tumor-infiltrating FOXP3⁺ Treg cells. Bar plots show the frequency of marker-positive cells within Tbet⁺ Treg and Tbet⁻ Treg subsets. Statistical significance was assessed using a paired t-test. Bar graphs represent mean ± SEM, and individual points indicate biological replicates; the square indicates SNC6, a cutaneous carcinoma with nasal extension.

A distinct cytotoxic CD4⁺ T cell cluster expressed canonical effector genes (*IFNG, GZMA*, *GZMB*, *PRF1*) and *PDCD1* (PD-1) (Fig. 3E). Cytotoxic CD4⁺ T cells have been described across human tumors and can acquire granzyme/perforin-mediated killing capacity while retaining checkpoint expression (Cachot et al. 2021; Patil et al. 2018; Quezada et al. 2010; Oh et al. 2020). Elevated *CXCL13* expression suggested a follicular helper T (Tfh) signature, but this cluster lacked additional canonical Tfh markers (e.g. *CD200*, *BCL6*), supporting its classification as cytotoxic rather than Tfh cells (Gu-Trantien et al. 2017).

Cells with a Treg signature dominated the intratumoral CD4⁺ T cell compartment and the four Treg subsets had a largely overlapping gene expression profile, with clear adaptations occurring between the Treg 1 to Treg 4 states (Fig. 3E, Supplementary Fig. 4A-B). All four Treg subsets (Treg 1-4) expressed high levels of *FOXP3*, *CTLA4* and *TIGIT* but features of tissue-resident, tumor-adapted Tregs, including *LAG3, CCR8, IL1R1* and *BATF* were more evident in the Treg 3 and 4 clusters (Fig. 3E, Supplementary Fig. 4B). CCR8⁺/BATF⁺ Tregs have been described as highly suppressive populations with stable residency in the TME (Plitas et al. 2016; Khatun et al. 2023). The Treg 3 and Treg 4 clusters were also enriched for *IL1R2* and *TNFRSF9* (4-1BB), markers associated with intratumoral Treg cell accumulation and suppressive function (Freeman et al. 2020; Dykema et al. 2023; Chen et al. 2022). In line with recent evidence distinguishing tumor-specific immune adaptations from generic inflammatory programs, these clusters also exhibited coordinated upregulation of *ICOS*, *IL1R1*, *IL18R1*, and *IL10*, resembling the ICOS⁺IL1R1⁺ subset previously described in human tumors (Mair et al. 2022). Notably, Treg 4 was enriched for *TBX21* (T-bet) and *CXCR3* (Supplementary Fig. 4B), consistent with a specific ability to dampen Th1-type inflammation and CD8⁺ T cell responses (Koch et al. 2009; Campbell and Koch 2011; Santegoets et al. 2019). These observations suggest that a substantial fraction of intratumoral Tregs in SNC adopt an activation-adapted, tissue-resident program, contributing to local immune suppression and functionally distinct from conventional inflammation-associated Tregs. A cluster of “dividing” cells expressing Treg cell-associated genes was also identified (Fig. 3E and Supplementary Fig. 4A), consistent with active proliferation of Treg cells within the TME.

Although solid tumors are often enriched for Treg cells, we compared the frequency of Treg cells among CD4⁺ T cells in SNC (as deciphered by scRNA-seq) to published datasets from squamous cell carcinoma of the head and neck (GSE173468), esophageal (GSE196756), non-small cell lung cancer (GSE198099), and normal tissue adjacent to head and neck tumors (GSE173468) (Supplementary Fig. 4C). SNC contained the highest frequency of cells with a Treg phenotype when compared with these conditions (Fig. 3F).

### T-bet, CXCR3 and 4-1BB are expressed by a portion of Treg cells in SNC

We next validated the major CD4⁺ T cell subsets by multiparametric flow cytometry in matched cryopreserved SNC and HNSCC tumors, corresponding peripheral blood samples, and healthy donor blood controls (Fig. 3G-K). Based on FOXP3 and TCF1 expression, CD4⁺ T cells segregated into three populations: Tregs (FOXP3⁺TCF1⁻), stem-like CD4⁺ (TCF1⁺FOXP3⁻) and a double-negative population (FOXP3⁻TCF1⁻). Both tumor types showed marked enrichment of FOXP3⁺ Tregs relative to blood, indicating tumor-specific accumulation, with Tregs accounting for on average, over 50% of intratumoral CD4⁺ T cells in SNC and ∼45% in HNSCC, despite substantial inter-patient variability (∼40% to >70%) (Fig. 3G).

Phenotypic profiling revealed that intratumoral Tregs expressed high levels of inhibitory and activation-associated receptors, including TIGIT, CTLA4 and 4-1BB (Fig. 3H-I), consistent with a chronically stimulated, suppressive phenotype and mirroring the scRNA-seq findings. The double-negative CD4⁺ T cell population contained cells expressing PD-1 and GZMB (Fig. 3H-I), and were enriched for cytotoxic CD4⁺ T cells. Conversely, the stem-like subset was enriched for IL-7R and CCR7 and lacked effector or inhibitory markers, consistent with a precursor CD4⁺ T cell state.

Although T-bet expression was highest in the cytotoxic CD4⁺ T cell subset, a substantial fraction of FOXP3⁺ Tregs expressed the transcription factor T-bet (Fig. 3J). Compared to their T-bet^-^ counterparts, a higher frequency of T-bet^+^ Treg expressed CXCR3, 4-1BB and PD-1 (Fig. 3K, Supplementary Fig. 4D), indicative of heightened activation, and suggesting that these may co-opt a Th1 priming environment for their adaptation. A minor FOXP3⁺CCR7⁺T-bet^-^ population was also detected (Fig. 3J), which may reflect the observation that cells in the stem-like 2 cluster expressed regulatory features including *FOXP3* and *IKZF2*. These data show that SNC tumors exhibit exceptionally high Treg infiltration, including CCR7⁺ and T-bet⁺ subsets, which is indicative of an environment that supports the differentiation and adaptation of Treg cells over time.

### Trajectory analysis of CD8⁺ T cells suggests two branches of differentiation

Although multiple T cell states were identified in SNC, their developmental relationships within the TME remained unclear. In particular, it is unknown whether stem-like, cytotoxic, TRM, and TExh cells represent divergent terminal fates or reflect continuous transcriptional adaptation. To address this, we applied pseudotime trajectory analysis to infer differentiation paths among intratumoral T cells based on continuous gene expression changes. Pseudotime revealed two principal trajectories originating from stem-like CD8⁺ T cells (Fig. 4A). One axis connected stem-like cells with TRM1/2 populations and extended toward TRM/Exh states, whereas a second axis progressed toward cytotoxic populations (Fig. 4A). The pre-TRM cluster occupied an intermediate position between these trajectories, consistent with a transitional state linking early differentiation with downstream effector or tissue-adapted programs.

**Figure 4.**
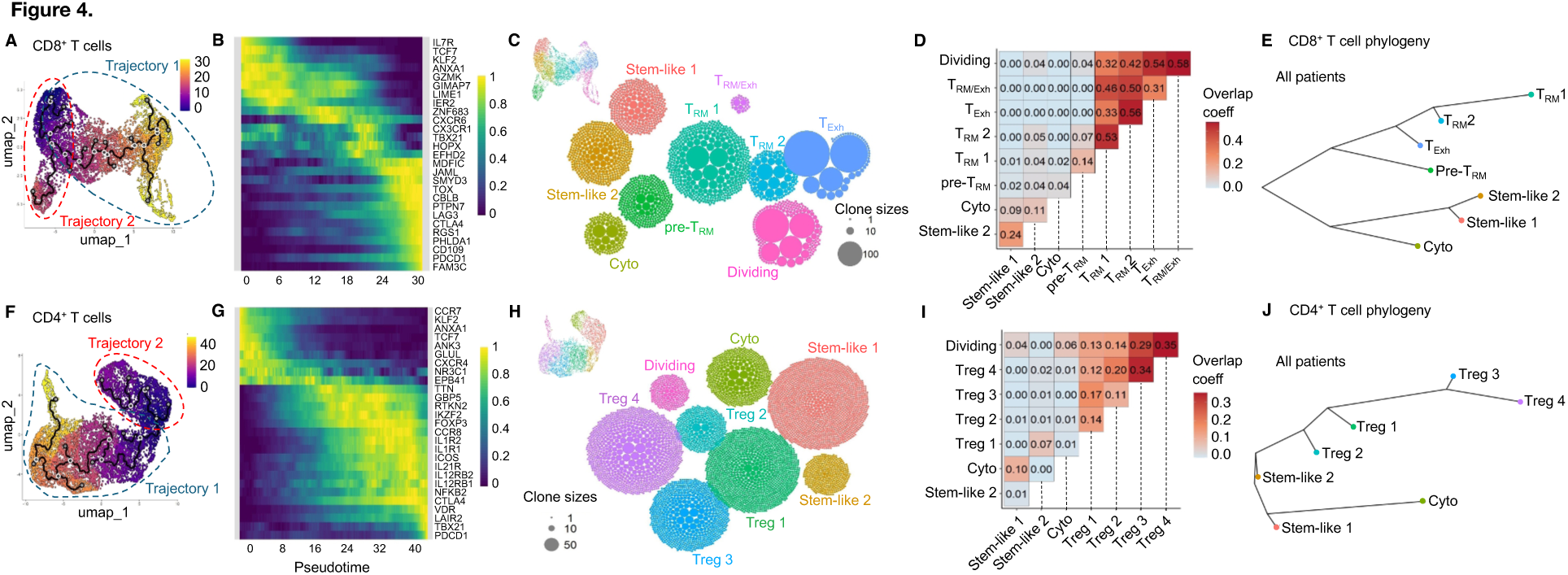
Pseudotime and PhyloTrajectory analysis deciphers differentiation pathways for CD4^+^ and CD8^+^ T cells in SNC. (**A**, **F**) Pseudotime trajectory analysis of intratumoral CD8⁺ (A) and CD4⁺ (F) T cells projected onto UMAP embeddings. Each dot represents a single cell, colored by pseudotime from early (dark purple) to terminal (yellow) states, inferred using Monocle 3. (**B**, **G**) Heatmaps showing relative expression of selected genes dynamically regulated along pseudotime for CD8⁺ (B) and CD4⁺ (G) T cells. Rows correspond to genes and columns represent ordered pseudotime bins. (**C**, **H**) Ball-packing plots illustrating clonal expansion across clusters for CD8⁺ (C) and CD4⁺ (H) T cells. Each ball represents a TCR clonotype contribution within a given cluster, with proportional size to the number of cells per clone in that cluster. Clusters are positioned and colored as in the UMAP. (**D**, **I**) Pairwise clonal overlap matrices for CD8⁺ (D) and CD4⁺ (I) T-cell states. Values indicate the overlap coefficient, reflecting the fraction of shared TCR clonotypes between pairs of clusters normalized to the smaller cluster size. (**E**, **J**) TCR-based phylogenetic trees inferred from clonotype frequency distributions across transcriptional states for CD8⁺ (E) and CD4⁺ (J) T cells from all SNC tumors. Branch topology and length reflect inferred lineage relationships and relative clonal flux between states.

Along the cytotoxic axis, cells progressively upregulated *CX3CR1* and *TBX21*, in line with a T-bet-driven effector program associated with peripheral migration and robust cytotoxicity (Kaech and Cui 2012; Gerlach et al. 2016) (Fig. 4B). Pseudotime-regulated gene expression further supported this bifurcation, revealing a continuum from IL7R⁺TCF7⁺ stem-like cells to either *ZNF683⁺* TRM states and *TOX*/*LAG3*-expressing exhausted cells, or toward TBX21⁺CX3CR1⁺ cytotoxic endpoints (Fig. 4B). These patterns define two dominant transcriptional programs within the CD8⁺ T cell compartment. Genes linked to transcriptional adaptation and restricted migration such as *RGS1*, *MDFIC* and *JAML* were enriched late in pseudotime (Bai et al. 2021; von Werdt et al. 2023; Oakley et al. 2017; Laukoetter et al. 2007). These transcriptional gradients are suggestive of a continuous but bifurcated CD8⁺ T cell differentiation spectrum within the TME (Miller et al. 2019; Kurtulus et al. 2019).

### PhyloTrajectory prediction supports bifurcated CD8⁺ T-cell differentiation in SNC

To test whether the transcriptionally inferred CD8⁺ T cell trajectories reflected true lineage relationships, we reconstructed clonal architecture using paired αβ TCR-V(D)J-seq data. Ball-packing visualization with *APackOfTheClones* (Ma et al. 2021) revealed extensive clonal expansions across all CD8⁺ subsets, particularly in the most differentiated states (TExh and TRM) (Fig. 4C). The dividing cluster also contained several large clones, consistent with ongoing antigen-driven proliferation (H. Li et al. 2019). Clonal diversity analysis supported these patterns, with cytotoxic and TRM/Exh clusters displaying high Gini coefficients and low Shannon diversity, indicative of oligoclonal expansion, whereas Stem-like 2 cells showed low Gini and high Shannon values, consistent with a highly polyclonal repertoire (Supplementary Fig. 5A-C).

Pairwise clonal overlap analysis revealed strong overlap in clonal repertoire between TRM and TExh clusters (correlation values > 0.3), relatively high overlap between the two stem-like clusters (corr. 0.24), and minimal clonal overlap between stem-like and more differentiated TRM and TExh states (Fig. 4D; Supplementary Fig. 5C). Notably, the transcriptionally intermediate pre-TRM cluster showed greater clonal relatedness to TRM subsets than to cytotoxic cells, suggesting its alignment within the exhaustion-associated trajectory rather than the cytotoxic branch.

We developed a probabilistic tool called PhyloTrajectory to infer T cell evolutionary relationships based on clonotype dynamics. PhyloTrajectory models changes in clonotype frequency along a phylogenetic tree as a continuous stochastic process (Methods). This approach allows sampling trees from posterior distribution that are compatible with the observed clonotype counts across the clusters. When applied to clonotype data across all patients, PhyloTrajectory revealed a shared branching trajectory between cytotoxic and stem-like (1 and 2) CD8⁺ T cell clusters, whereas TRM and TExh clusters segregated along a distinct branch associated with pre-TRM cells (Fig. 4E). These results are consistent with pseudotime analysis but also suggests that TRM and TExh states may be self-sustaining or replenished in part by pre-TRM cells, while the relatively large pool of stem-like CD8^+^ T cells showed limited clonal relatedness to TRM and TExh in the TME and instead branched primarily with the cytotoxic CD8^+^ T cell cluster.

### Trajectory analysis of CD4⁺ T-cells suggests separate clonal branches for cytotoxic and Treg cells

Pseudotime inference revealed two main differentiation trajectories among intratumoral CD4⁺ T cells (Fig. 4F). The dominant pathway originated from stem-like populations and progressed sequentially through Treg 1-4 states. The second trajectory branched toward the cytotoxic CD4⁺ subset. Along the Treg trajectory, early stem-like cells expressing *CCR7*, *TCF7* and *KLF2* progressively acquired a regulatory transcriptional program, with gradual upregulation of *FOXP3*, *IKZF2*, *CTLA4* and *ICOS*, together with cytokine receptor components *IL12RB1* and *IL12RB2*. Late pseudotime was characterized by the induction of tumor-resident genes and highly suppressive Treg states, including *CCR8*, *VDR* and *LAIR2*, together with inflammatory sensing and regulatory receptors such as *IL1R1* and *IL1R2*. In parallel, transcriptional regulators linked to activation and type 1-associated programs, including *NFKB2* and *TBX21*, were upregulated, consistent with the emergence of a highly differentiated T-bet⁺ Treg population (Fig. 4G).

### PhyloTrajectory suggests that clonal evolution of CD4^+^ T-cells in SNC occurs along two primary branches

In CD4⁺ T cells, clonal expansion was evident in the most differentiated subsets, particularly the Treg 3, Treg 4 and cytotoxic clusters (Fig. 4H). Clonal diversity metrics revealed broader repertoire diversity in CD4⁺ than CD8⁺ T cells, with higher Chao1 values reflecting extensive clonal richness, consistent with the prevalence of single clones in the CD4⁺ compartment (Supplementary Fig. 5D-F). The Treg clusters were the most clonally related, in particular the Treg 3 and Treg 4 clusters (correlation 0.34) (Fig. 4I). The dividing population shared clones with both Treg and Cytotoxic CD4^+^ T cell clusters, indicating proliferation along both differentiation trajectories.

The PhyloTrajectory model placed stem-like 1 and 2 clusters near the base of the tree, a feature of their poor clonal expansion. Cytotoxic CD4⁺ T cells had a long branch, indicative of substantial clonal turnover that was distinct from that of Treg cells. The most expanded populations, Treg 3 and Treg 4 cells, shared a long branch with Treg 1 and Treg 2 states. Thus, CD4 T cells in SNC appeared to differentiate along two primary branches, one supporting the cytotoxic state, and another supporting the Treg cell states.

### The ‘Stem-like’ compartment is comprised of cells primed for Treg cell differentiation

We observed that some cells in the stem-like clusters expressed *FOXP3* and *IKZF2* mRNA (Fig. 3E), and that approximately 10% of gated FOXP3^+^ Treg cells expressed CCR7 protein (Fig. 3I-J) by flow cytometry analysis. Some clonal overlap was detected between the Treg clusters and cells in the stem-like clusters (Supplementary Fig. 5F-G). We next analyzed the phenotype of stem-like CD4^+^ cells with a verified clonal relationship to cells of *bona fide* Treg clusters (Treg 1-4). Among stem-like clones that were shared with cells in the Treg clusters (36 distinct clones in total across patients), approximately 60% expressed *FOXP3* mRNA at the stem-like stage (Supplementary Fig. 5H-I). These stem-like clones also expressed *IL7R, CCR7, LEF1, ANXA1*, and other stem-like features (Supplementary Fig. 5H-I). Thus, the CD4^+^ stem-like compartment appears to include cells that are primed towards the Treg cell fate.

Together, pseudotime and PhyloTrajectory analyses support differentiation of CD4⁺ and CD8⁺ T cells along multiple, clonally distinct pathways in SNC.

### Spatial V(D)J profiling identifies T and B cells in close proximity in SNC

Single-cell sequencing technologies allow us to decipher thousands of paired αβTCR and IGH/L clones. However, a limitation of these approaches remains the ability to directly visualize lymphocyte clones in the tissue in which they reside. To resolve spatial locations of specific B and T cell clones, we employed the spatial V(D)J approach we recently described (Engblom et al. 2023) (Fig. 5A). Starting from amplified, spatially barcoded cDNA generated on the Visium platform, we performed a targeted enrichment of BCR and TCR sequences (including γδ chains), followed by PacBio HiFi library preparation and sequencing. Importantly, this enrichment is not clone-specific and enables broad capture of TCR and BCR transcripts without prior selection of individual receptor sequences. This enabled recovery of full-length receptor sequences spanning the complete V(D)J and constant segments, spatial barcode and UMI (Fig. 5A). Hundreds of unique TCR sequences were detected across the three patient tissues, with SNC4 displaying the highest apparent clonal diversity, with 702 unique TCRβ (*TRB*) sequences. Notably, the spatial V(D)J approach does not capture paired TRA:TRB or TRG:TRD sequences. Consequently, spatial clonotypes, or clones, are defined based on the unique sequences of individual receptor chains. TRB chains comprised the majority of clonotypes detected in each patient (Fig. 5B), in line with reports showing that TCR sequencing methods typically capture TRA diversity less efficiently (Barennes et al. 2021). TRG and TRD chains were detected at low frequencies in SNC2 and SNC4 and were not detected in SNC3. To assess whether TCRβ sequences alone could approximate clonal identity, we quantified the extent to which identical TRB CDR3 nucleotide sequences were associated with multiple clonotypes (defined by paired TRA and TRB chains) in the single-cell dataset. We found that across both CD4⁺ and CD8⁺ T cells, the vast majority of TRB sequences uniquely mapped to a single clonotype (>96%), with only a small fraction associated with multiple clonotypes due to alternative TRA pairings (Supplementary Fig. 6A). Notably, TRB sequences associated with multiple clonotypes were typically low-frequency events, suggesting a limited impact on downstream analyses. We then compared the abundance of spatial clones to those recovered from single-cell analysis on a per patient basis. Clone abundance was correlated across both platforms (Fig. 5C) suggesting that the pieces of tissue used for single-cell dissociation and for spatial analysis were not highly divergent in T cell composition.

**Figure 5.**
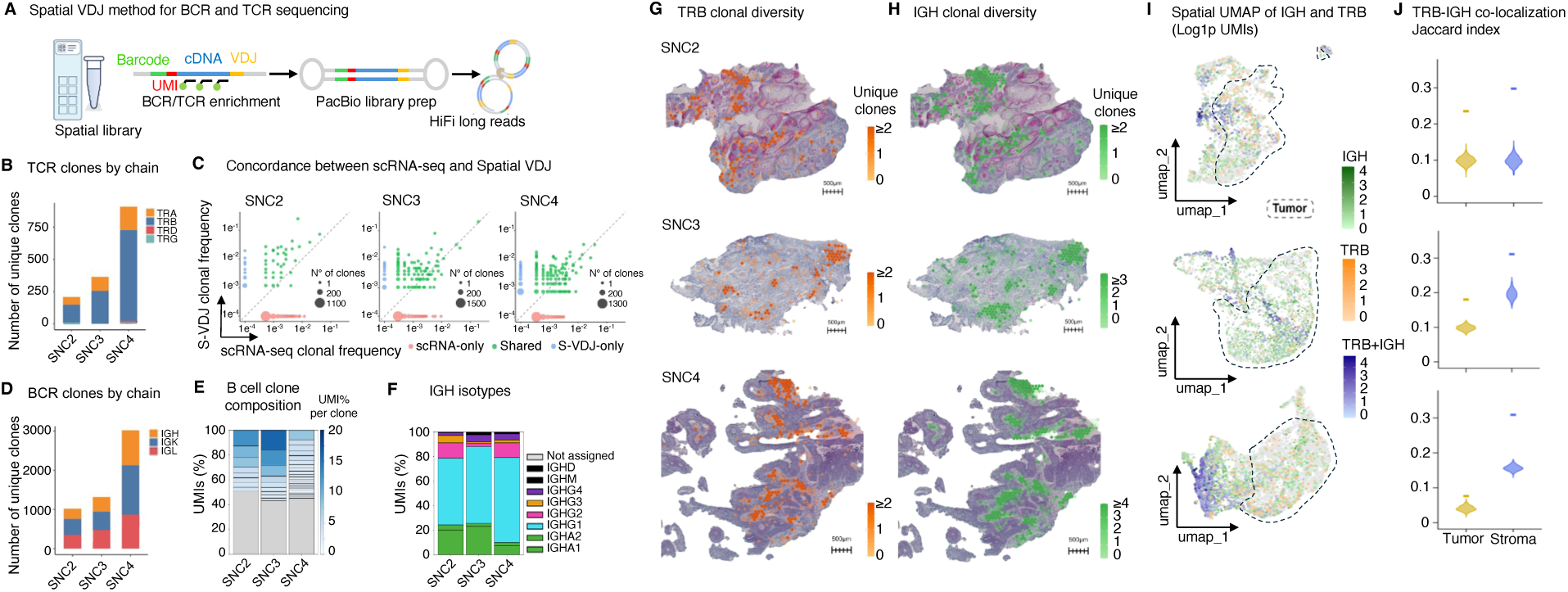
Spatial V(D)J reveals frequent colocalization of T and B cells in SNC. (**A**) Schematic overview of the spatial V(D)J sequencing workflow. Visium-derived amplified cDNA undergoes targeted enrichment of BCR and TCR transcripts followed by long-read sequencing, enabling recovery of full-length immune receptor sequences while preserving spatial barcode information. (**B**) Number of unique TCR sequences recovered (TRA, TRB, TRG, TRD) from spatial tissue sections across patients. (**C**) Scatter plots showing clonotype frequencies detected by scRNA-seq (x-axis) and spatial transcriptomics (y-axis) across samples. Shared clonotypes are in green. Modality-specific clonotypes are in red (scRNA-seq) or blue (spatial). Point size reflects clonotype abundance and the diagonal line indicates equal relative frequencies across modalities. (D) Number of unique BCR sequences recovered (IGH, IGK, IGL) from spatial tissue sections across patients. (**E**) Bar graph of B cell clone frequency. Top 5% of clones are segmented while the 95% least abundant clones are shown in gray. (**F**) IGH isotype distribution across spatial tumor sections, shown as the fraction of UMIs. (**G, H**) Spatial distribution of unique TRB (G) and IGH (H) clonotypes, mapped onto a representative SNC2-4 tumor section. IGH UMI threshold >1. Scale bars, 500 µm. (**I**) Spatial transcriptomic UMAPs showing log-transformed TRB and IGH UMIs per spot for each patient. Spots are classified as TRB-only (orange), IGH-only (green), dual IGH^+^TRB^+^ (blue), or none (grey). (**J**) Overlap between TRB^+^ and IGH^+^ spots was quantified separately in tumor and stromal compartments for each patient using the Jaccard index. Violin plots show the permutation-derived null distribution generated by randomly shuffling TRB-positive spots within each compartment, and horizontal colored lines show the observed Jaccard index. Tumor and stromal compartments were defined based on merged spatial clusters in Fig. 1F.

Next, we analyzed the clonal identity of IGH, IGK, and IGL chains captured by Spatial V(D)J. Several hundred IGH clones were found in each patient, with SNC4 again exhibiting the highest abundance of unique BCRs (Fig. 5D). We removed BCRs with only a single UMI and analyzed clonal abundance across the 3 patient tumors. The top 5% of clones contributed approximately 50% of all BCR UMIs, indicating that some B cell clones were highly expanded (Fig. 5E). BCRs of an IgG subclass were most abundant across all three tumors, although a considerable proportion of UMIs also belonged to the IgA subclass, in line with the mucosal function of the sinuses (Fig. 5F).

To visualize T and B cell clonal diversity in tumor tissue, the number of unique TRB and IGH clonotypes per spot was mapped spatially across the three tumors (Fig. 5G-H) and on the spatial RNA-Seq UMAP space (Fig. 5I). TRB and IGH clonotypes showed spatially restricted distributions, forming discrete immune-rich niches that showed some overlap within each tumor (Fig. 5G-I). Consistent with this, permutation-based Jaccard analysis showed that TRB-positive and IGH-positive spots co-localized more than expected by chance in both tumor and stromal compartments across all three tumors (Fig. 5J), supporting the presence of shared adaptive immune niches within the TME.

### Single T and B cell clones are highly dispersed throughout the TME

Because the TRB CDR3 sequence uniquely identified shared clones between our scRNA-seq dataset and the tissue section, it was possible to map CD4⁺ and CD8⁺ T cell clones directly in the tumor tissue and infer their transcriptional phenotypes (Fig. 6A-B, Supplementary Fig. 6B). Spatial mapping revealed that expanded CD8⁺ T cell clones were predominantly associated with TRM and TExh phenotypes. Visualization of representative expanded CD8⁺ clonotypes revealed broad spatial dispersion of single clones across multiple regions in all three tumors. Expanded CD4⁺ T cell clones were likewise distributed across the tumor tissue and were mainly linked to Treg 1-4 states. Consistent with their smaller clonal size, CD4⁺ clonotypes appeared less abundant than their CD8⁺ counterparts (Fig. 6A-B).

**Figure 6.**
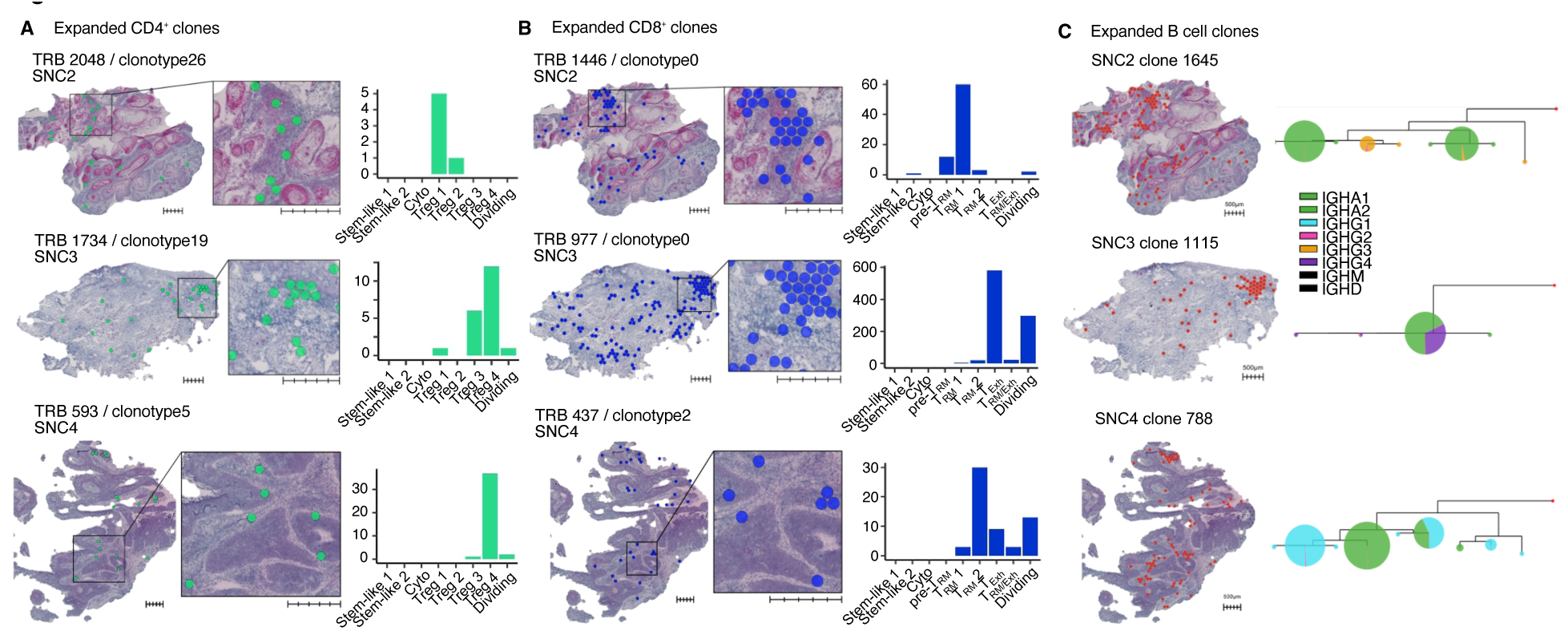
Precise clone mapping identifies Treg clones in close proximity to TRM, TExh and activated B cells in SNC. (**A, B**) Representative expanded CD4⁺ (A) and CD8⁺ (B) T cell clonotypes in SNC2, SNC3 and SNC4. For each sample, the left panel shows the spatial distribution of the indicated clontype and the inset shows a zoom-in of the boxed region. All scalebars, 500 µm. Bar plots (right) show the cluster distribution of the indicated clonotype. (**C**) Spatial mapping of representative expanded B cell clonotypes onto tumor sections (left). Phylogenetic reconstruction of clonotypes from each SNC tumor (right). Circle size reflects sequence abundance within clone, and colors indicate isotype identity.

We next examined the IGH repertoires of the three SNC patients from our spatial V(D)J analysis. Clones resolved by spatial V(D)J in all patients showed varying degrees of somatic hypermutation (SHM) (Supplementary Fig. 6C). Representative single expanded B cell clones appeared to be dispersed around the tumor and undergoing clonal evolution in SNC, as highlighted by their complex tree structure and evidence of class switching (Fig. 6C). Analysis of the single-cell transcriptomes of B cells in SNC revealed cells of a predominantly activated phenotype (CD27, TNFRSF13B, IFN-induced genes), and a small population of plasma cells (Supplementary Fig. 6D-F). Activated B cells did not express canonical tertiary lymphoid structure (TLS) markers such as *BCL6*, *CXCL13* and *AICDA* (Supplementary Fig. 6D-F). Flow cytometric analysis revealed that B cells expressed *CXCR3* and displayed a tissue-resident phenotype (CD69⁺) in SNC indicating that these cells may be following similar chemotactic gradients as T cells (Supplementary Fig. 6G). Thus, clone tracking identifies a highly dispersed distribution of single T and B cell clones across the SNC TME.

### T and B cells are localized in niches enriched for genes associated with tissue stroma and antigen presentation

To determine whether Treg, CD8^+^ T and B cells inhabited specific repetitive niches in SNC, we applied non-negative matrix factorization (NMF) to a weighted gene-by-spot matrix and quantified the correlation between each factor with the number of unique TRB or IGH clones per spot. This identified shared transcriptional programs associated with clonal diversity across patients (Fig. 7A). Notably, the three NMF factors showing the strongest positive correlations with TRB diversity (NMF_10, NMF_16 and NMF_17) were also among the strongest correlates of IGH diversity, suggesting convergence of B and T cell diversification within shared spatial niches. Based on their top-weighted genes, NMF-17 was enriched for plasma cell markers, confirming our observations that TRB and IGH densities were spatially linked (Fig. 7A, Supplementary Fig. 7A). NMF_10 represented a pronounced desmoplastic, fibroblast-rich stromal program with top-weighted features including collagens (*COL1A1*, *COL1A2*, *COL3A1*), extracellular matrix proteins (*SPARC*, *DCN*, *BGN*, *LUM)* and cancer-associated fibroblast markers (*COL6A2, AEBP1*, *POSTN*, *IGFBP7*) (Supplementary Fig. 7A). NMF_16 was enriched for MHC-II genes (*HLA*-*DRA/DRB1/DQB1/DPA1/DPB1)*, *CD74*, chemokines (*CXCL9*, *CXCL10*), and interferon-induced transcripts (*IFI30, WARS*, *TAP1, ISG15*), defining an IFN-activated antigen-presenting cell (APC) program. We also detected expression of *C1QA* and *C1QB,* indicating the presence of phagocytic C1Q⁺ macrophages at these sites (Supplementary Fig. 7A). Together, these findings suggest that T and B cells are closely associated with CAFs and APCs within the TME.

**Figure 7.**
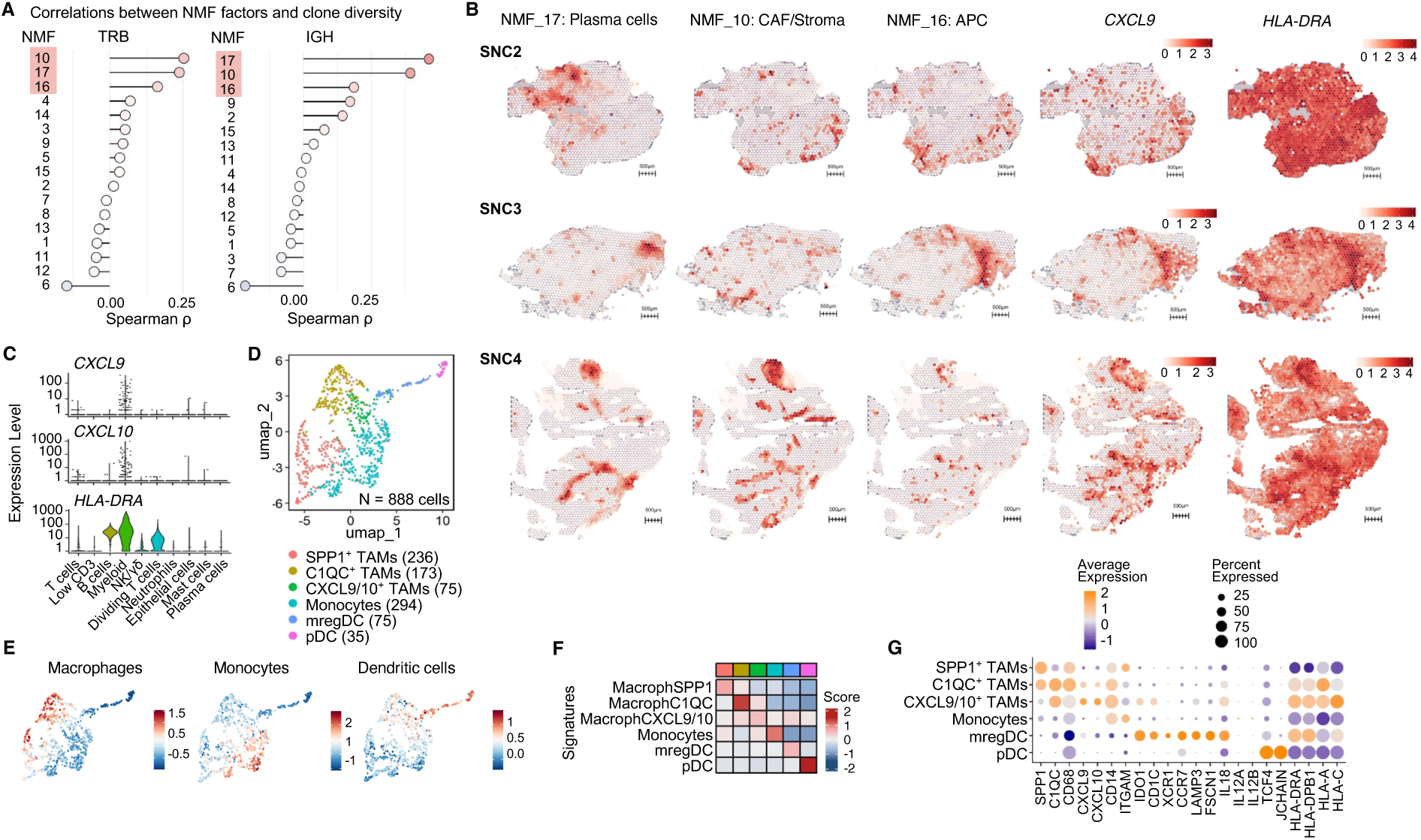
T and B cell clonal diversity is associated with CAFs and APCs in SNC. (**A**) Ranked Spearman correlations between NMF factor activities and spatial TRB (left) or IGH (right) clonotype diversity across all Visium spots. The 3 top factors are highlighted in red. (**B**) Spatial maps of selected NMF factors, CXCL9, and HLA-DRA expression. In gene expression maps, color scale values indicate UMI counts, whereas in NMF maps, the scale indicates factor activity. Scale bars, 500 µm. (**C**) Violin plots of *CXCL9*, *CXCL10*, *IDO1*, and *HLA-DRA* expression across cell types in the scRNA-seq dataset. (**D**) UMAP of tumor-infiltrating myeloid cells resolved into six subsets: SPP1⁺ tumor-associated macrophages (SPP1⁺ TAMs), C1QC⁺ TAMs, CXCL9/10⁺ TAMs, monocytes, mregDC, and plasmacytoid dendritic cells (pDC). (**E**) UMAP projections of curated macrophage, monocyte, and DC gene signatures. (**F**) Heatmap of average module scores for these signatures across myeloid subsets. (**G**) Dot plot showing the average expression and percent of cells expressing selected genes across myeloid subsets.

We analyzed factor activity side-by-side across tumors spatially (Fig. 7B). Regions of high factor activity were dispersed across the tumor in line with the distribution of T and B cell clones. Some tumor regions were rich for all factors at once (particularly evident in SNC4) while other regions showed preferential enrichment for specific factors (Fig. 7B). Several genes associated with NMF_16 including *CXCL9*, *CXCL10* and *HLA-DRA* were robustly detected at discrete sites in each tumor section (Fig. 7B, Supplementary Fig. 7B).

### SNC antigen presenting niches are made up of CXCL9/10^+^ macrophages and CCR7^+^IDO1^+^ dendritic cells

To explore the cell populations contributing to the NMF_16 APC factor, *CXCL9*, *CXCL10* and *HLA-DRA* expression was analyzed in our scRNA-seq dataset. These genes were largely confined to the myeloid cluster, with some expression of HLA-DRA in the T and B cell clusters (Fig. 7C). Re-clustering of intratumoral myeloid cells identified tumor-associated macrophage (TAM), monocyte and dendritic cell (DC) populations, including a subset of DC enriched for markers of ‘mregDC’, recently implicated in suppressing immunity in solid tumors (Maier et al. 2020) (Fig. 7D-G).

Among macrophages, three subtypes were identified including CXCL9/10^+^, C1QC⁺ and SPP1^+^ TAMs. C1QC⁺ TAMs exhibited phagocytic programs and expressed moderate levels of HLA class II genes (Zilionis et al. 2019). SPP1⁺ TAMs expressed inflammatory and angiogenic mediators previously associated with tumor progression across multiple cancers (Bill et al. 2023; X. Li et al. 2025). CXCL9/10⁺ TAMs displayed high levels of *HLA* class I and II (Fig. 7G), suggestive of a capacity to affect anti-tumor immunity via recruitment of CXCR3-expressing lymphocytes (House et al. 2020). Importantly, SPP1⁺ TAMs clustered within tumor regions and were segregated from CXCL9/10^+^ regions (Supplementary Fig. 7B), in line with recent findings (Bill et al. 2023). Overall, these data indicate that a subset of TAMs expressing CXCL9/10 are associated with the presence of T and B cells.

Although DC are known to express CXCL9 and CXCL10 in lymphoid tissues and prime Th1 and CD8⁺ T cell responses (Groom et al. 2012), they did not express these chemokines in SNC. Instead, tumor-infiltrating DC exhibited an inflammatory/regulatory phenotype characterized by *IL18*, *CCR7, LAMP3*, *FSCN1* and high *IDO1* expression, an enzyme associated with the induction and maintenance of Treg cells (Munn and Mellor 2007; Holmgaard et al. 2013)*. IDO1 mRNA* expression overlapped with *CXCL9* and *CXCL10* expression in SNC tumor sections (Supplementary Fig. 7B), indicating that the tumor APC niche could be comprised of both CXCL9/10⁺ TAMs and CCR7^+^IDO1^+^ DC.

### Fluorescence microscopy validates the existence of complex peritumoral immune niches in SNC

To validate the spatial localization of T and B cells in distinct niches in SNC, we performed confocal microscopy on cryopreserved tumor sections. Imaging revealed large PanCK^+^ tumor epithelium regions surrounded by dense infiltrates of CD3^+^ T cells (Fig. 8A, Supplementary Fig. 8A). In several cases, T cells preferentially localized to PDPN⁺ stromal regions consistent with fibroblast-rich areas (Fig. 8A), although PDPN expression was occasionally also detected in tumor epithelium (not shown), in line with reports indicating PDPN expression on squamous cell carcinomas may be a marker of poor prognosis (Hesse et al. 2016; Krishnan et al. 2018). In depth analysis of CD3-rich zones showed frequent nuclear FOXP3 expression in T cells (Fig. 8B-D), and a close proximity of CD4^+^ and Treg cells to B cells (Fig. 8B) and to IDO1^+^LAMP3^+^ DC (Fig. 8C). CXCR3 expression was detected in both CD8^+^ T cells and Treg cells in these zones (Fig. 8D), while CXCL9-expressing cells were found in close proximity to T cells (Fig. 8E), consistent with a chemokine-rich microenvironment supporting immune cell recruitment and positioning.

**Figure 8.**
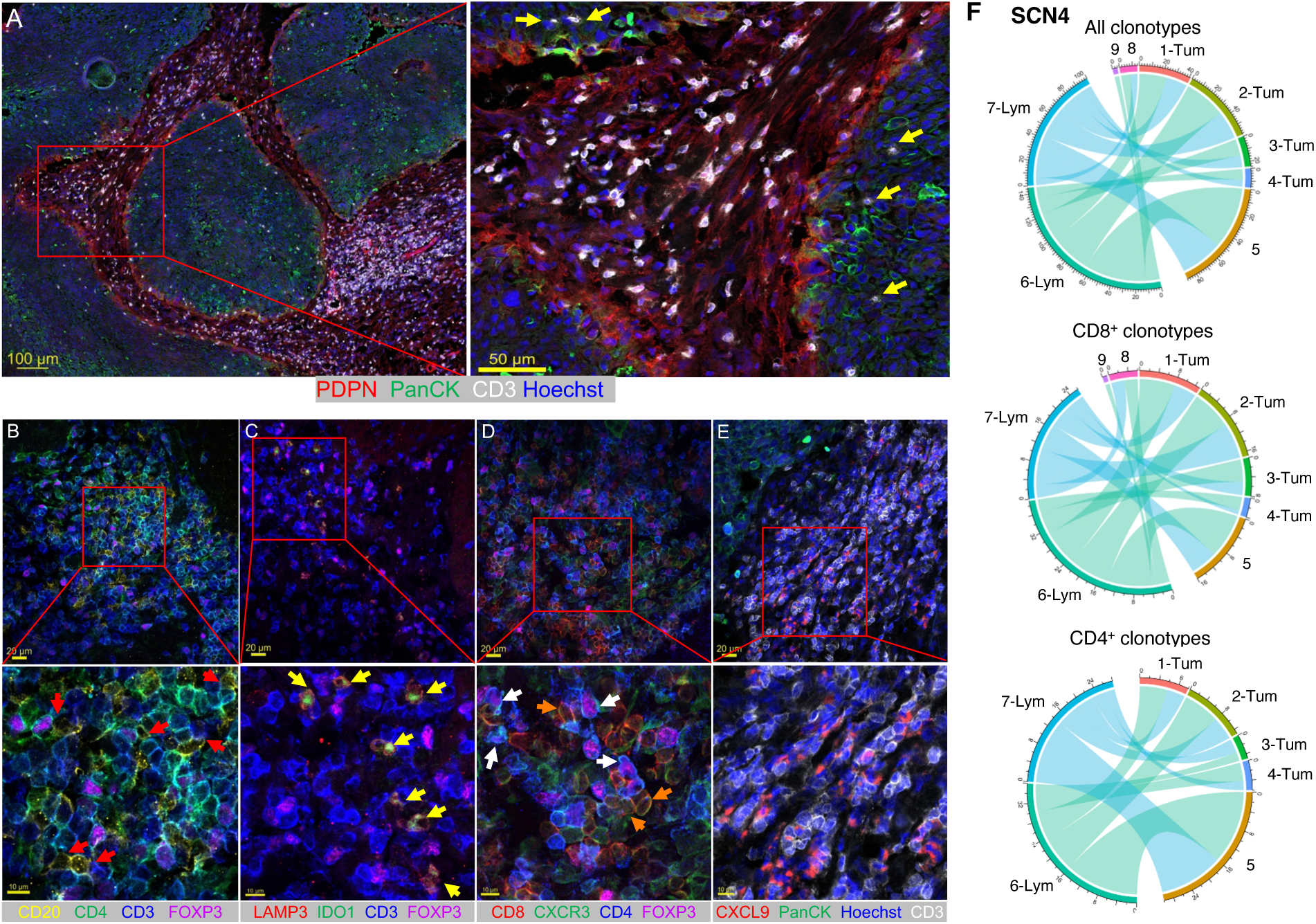
T cells inhabit complex multicellular niches and are clonally related to tumor epithelium-invading T cells. (**A**) Confocal images of SNC tumor sections showing PanCK⁺ tumor epithelium (green), PDPN⁺ stromal regions (red), CD3⁺ T cells (white), and nuclei (Hoechst, blue). Left, overview of tumor architecture; right, magnified view highlighting dense CD3⁺ T cell infiltration in PDPN⁺ stromal regions and in tumor epithelium regions (yellow arrows). Scale bars, 100 μm (left) and 50 μm (right). (**B**-**E**) Higher-resolution images of immune cell interactions within CD3-rich regions. Upper panels show broader views (scale bars, 20 μm), with corresponding magnified regions below (scale bars, 10 μm). Markers and colors are indicated. (**F**) Chord diagram of SNC4. Clones from lymphocyte-rich regions (clusters 6 and 7) that were present in other spatial clusters were plotted. Shown are ‘all clonotypes’ as aligned by spatial V(D)J, and specific CD8^+^ and CD4^+^ clones, which could be verified by scRNA-seq.

### T cells in the peritumoral space are clonally-related to those invading tumor epithelium regions

Although T cells were enriched in peritumoral regions, they also visibly infiltrated the PanCK^+^ tumor epithelium regions (Fig. 8A, yellow arrows, Supplementary Fig. 8A-B). To directly assess whether lymphocyte-rich regions may be supporting T cell invasion of the tumor epithelium regions, we analyzed the clonal relationships between these regions. T cell clones in peritumoral regions were clonally-related to those infiltrating the tumor (Fig. 8F, Supplementary Fig. 8C-D). Both CD4^+^ (primarily Treg cells) and CD8^+^ clones were observed to penetrate the tumor epithelium space across all tumors (Fig. 8F, Supplementary Fig. 8C-D), demonstrating that peritumoral regions supported T cell populations with the potential to infiltrate tumor epithelium regions.

Taken together, these results suggest that SNC is composed of evolving T and B cell populations, which are shaped by complex interactions between tumor epithelium, CAFs, and antigen presenting cells. Although peritumoral CXCL9-rich niches appear to support profound T cell accumulation, the high frequency of Treg cells and presence of IDO1^+^ DC within these niches may impede anti-tumor immunity. These findings have implications for the treatment of SNC and more common HNSCCs.

## Discussion

In this study, we used scRNA-seq, spatial transcriptomics and untargeted spatial V(D)J profiling to delineate the architecture of SNC and place T and B cell clones and subsets within their spatial context.

The integration of detailed single-cell phenotyping with spatial information has been challenging, particularly for the highly heterogeneous T cell compartment. Recent studies have addressed this by using scRNA-seq or bulk RNA-seq to identify dominant TCR clonotypes, followed by targeted spatial profiling using platforms such as Xenium or Cartana *in situ* sequencing (McCord et al. 2026, Yu et al. 2025).

In contrast, although our approach does not achieve single-cell spatial resolution, it enables untargeted, large-scale spatial profiling of TCR and BCR repertoires directly within intact tumor tissue. This allows detection of a broader spectrum of adaptive immune clones, including low-frequency or spatially segregated clonotypes that may be missed by targeted approaches, while simultaneously enabling parallel analysis of T and B cell clonality. Together, these results demonstrate that untargeted spatial V(D)J profiling provides a complementary and scalable strategy for studying adaptive immune organization in human tumors, offering insights into clonal diversity and spatial immune architecture. In addition, we expanded our pipeline to detect γδ T cells, which were only sparsely present in SNC, suggesting a limited role for these cells.

An additional advantage of spatial V(D)J profiling is its ability to directly map T- and B-cell clonotypes within tumor tissue. Lymphocyte markers such as CD3 chains, CD79A, and CD20 can be sparse and diluted on the Visium platform, meaning that localization based on gene expression alone may be less sensitive or specific in some regions. Spatial V(D)J profiling enabled clone-resolved mapping, allowing confident spatial association with CAFs and IFN-exposed APC niches. The observation that identical T or B cell clones were detected across multiple spatially distinct lymphoid hubs within individual tumors indicates a degree of intratumoral lymphocyte migration, potentially via the vasculature. This is consistent with recent spatial TCR profiling in head and neck cancer, where presumed tumor-specific T cells and tumor-enriched clones were broadly distributed across the tumor microenvironment (McCord et al. 2026). Together, these findings suggest that distinct lymphoid foci, supported by underlying stromal or APC compartments, constitute the primary microenvironments in which lymphocytes accumulate, persist and clonally expand, in close proximity to the tumor epithelium. It should be noted that although lymphocytes were predominantly localized within large hubs segregated from the tumor epithelium, sparse but recurrent T cells were also found infiltrating the tumor epithelium compartment, a feature that has been associated with improved prognostic outcomes (Liang et al. 2025). In a translational setting, this architecture-preserving clonal readout could help stratify solid cancer patients and monitor how therapies reshape local immune niches.

Our linked single cell and spatial analysis revealed many interesting immunological features of SNC, a subtype of HNSCC that has not been extensively studied. One prominent feature was the exceptionally high frequency of FOXP3^+^ Treg cells. Although solid tumors are known to be enriched for Treg cells (Togashi et al. 2019; Plitas and Rudensky 2020), frequencies in SNC often exceeded 50% of CD4⁺ T cells, which is among the highest of all solid cancers. How Treg cells come to reside in SNC at such high frequencies may depend on several factors. Firstly, Treg cells expressed the vitamin D receptor, which can upregulate FOXP3 expression in human T cells (Matos et al. 2022; L Bishop et al. 2020). The close proximity of Treg cells to IDO1^+^ DCs, known to promote Treg stability, likely further reinforces local immune tolerance (Munn and Mellor 2007; Holmgaard et al. 2013). It appeared that the SNC TME supported Treg cell differentiation from the stem-like CD4^+^ T cell state to a fully tumor-adapted IL1R1^+^CCR8^+^ state. Although *FOXP3* mRNA was already detected in the stem-like CD4^+^ T cell compartment, clone tracking suggested that SNC Treg may also differentiate from FOXP3^-^ stem-like CD4^+^ T cells, often dubbed ‘inducible’ Treg cells. The robust detection of *FOXP3* among dividing cells was also indicative of a robust environment for Treg cell differentiation and expansion. These observations suggest that Treg cells play an important role in regulating tumor immunity in SNC.

Some modes of Treg cell-mediated immune regulation seem most plausible, based on the proximity of Treg cells to CD8^+^ T cells and IDO1^+^ DC in the SNC TME. Strong T-bet and CXCR3 expression is indicative of a TME that selectively restrains IFN-γ-producing CD8⁺ and CD4⁺ Th1 cell responses (Koch et al. 2009; Levine et al. 2017; Moreno Ayala et al. 2023). Studies in mice have clearly shown that T-bet^+^CXCR3^+^ Treg cells impair anti-tumor immunity (Moreno Ayala et al. 2023), but this has typically been attributed to the ability of Treg cells to restrain CD8+ T cell priming in draining lymph nodes. The close proximity of Treg cells with CD8 T cells in SNC suggests that such a mechanism of regulation may occur directly in the SNC TME. Another possibility is that Treg cells dampen the function of tumor-infiltrating DC, as shown by a recent study where Treg cells reduced CD40 expression on CCR7^+^ DC, thereby limiting CD4^+^ and CD8^+^ T cell activation (Zitti et al. 2026). Clonally-expanded Treg cells in SNC were also most capable of IL-10 production, which is known to restrain immunity by several mechanisms. Thus, targeting Treg cells in SNC may prove highly efficacious, even if Treg cell-specific therapies are yet to translate from preclinical to clinical settings.

The transcriptomic and clonotypic features of the SNC CD8^+^ T cell population were also of interest. SNC CD8^+^ T cells could be segregated into TRM and TExh populations on a transcriptional basis. Specifically, TRM 1 had a low exhaustion score, while vice versa TExh cells had a low resident memory score. Our annotated TRM 2 cluster did share transcriptional features of both resident memory and exhausted cells, in line with a recent study of breast cancer, where traditional gene signatures could not discern TRM from TExh cells (Burn et al. 2026). In breast cancer, refined signatures were able to parse out TRM and TExh cells, which were shown to be largely clonally-distinct, with TExh cells associated with tumor antigen-specific responses and TRM cells related to clones from normal healthy breast tissue. This is a point-of-difference with SNC, where TRM and TExh cells appeared to differentiate along the same clonal trajectory. Our novel PhyloTrajectory tool deciphered clonal relationships between tumor stem-like, TRM and TExh cells that have been difficult to discern previously. Although stem-like CD8^+^ T cells from lymph nodes are known to support the tumor-infiltrating TRM and TExh cell populations (Wijesinghe et al. 2025), it has not been clear if stem-like CD8^+^ T cells in the tumor tissue served the same function. Our results suggest that stem-like CD8^+^ T cells in SNC tumors share a clonal trajectory with cytotoxic cells, but not with TRM and TExh cells in SNC. This indicates that the TRM and TExh populations may be in part sustained by cells from the pre-TRM cluster, in particular in SNC, where lymph node involvement is rarer than in other solid cancers.

In all, the underlying tumor architecture likely plays a critical role in patient outcomes and response to immune therapies, yet this aspect is often overlooked. Technological improvements now enable clonal and spatial niche analyses directly within tumor tissue, which reveals that the SNC TME supports extensive adaptation of T and B lymphocytes in juxtaposition with innate immune cells, CAFs and the tumor itself. As studies continue to delineate the architecture of a diverse range of solid tumors, we anticipate that patient prognosis and the design and use of immune-based therapeutics will become more efficacious and targeted.

## METHODS

### Study Design

All procedures performed in this study involving human samples were approved by the Swedish Ethical Review Authority and followed the ethical standards and guidelines of the institutional and/or national research committee and the Declaration of Helsinki. All patients were treated according to the standard treatment protocol at Karolinska University Hospital, Stockholm, Sweden. Informed and signed consent was obtained from all individual participants included in the study. This study involved human participants and was approved by the Ethics Committee (2019-03518 and 2021-01265). Data was processed in accordance with applicable data-protection regulations. Patients were assessed and determined to have sinonasal cancers of squamous cell origin and were thereafter scheduled for surgery at Karolinska Hospital, Stockholm, Sweden. Patient tumors were resected and were transferred to researchers on the same day.

### SNC tumor dissociation and cell preparation for scRNA-seq

Fresh tumor samples from SNC patients were dissociated by chopping with scissors, followed by gentle trituration through a 70 μm filter using Iscove’s Modified Dulbecco’s Medium (IMDM) containing 10% heat-inactivated FBS and 1% penicillin/streptomycin. No enzymatic digestion was performed. Red blood cell lysis was carried out using Invitrogen™ eBioscience™ 10X RBC Lysis Buffer prior to staining. For analysis with the Chromium (10x Genomics) platform, cell purification was performed by flow cytometry with a FACSAria Fusion (BD Biosciences) with a preference for CD3^+^ T cells, but also including a portion of non-T cells before loading onto the 10x Chromium platform. Excess tumor material or matched patient blood was cryopreserved for further analysis.

### Spatial transcriptomics

#### Sample Collection

Sinonasal tissue samples were obtained from 3 patients (SNC2-4, Supplementary table 1). The tumor samples were immediately snap-frozen in OCT on a chilled metal plate over dry ice to preserve the RNA integrity. The frozen tissue sections were stored at -80°C until further processing.

### RNA extraction and quality control

Total RNA was extracted from cryosectioned tumor tissue using the RNeasy Plus Mini Kit (Qiagen) according to the manufacturer’s instructions with minor modifications. Briefly, six 10-µm cryosections per sample were homogenized in RLT Plus buffer supplemented with β-mercaptoethanol using bead-based mechanical disruption. RNA integrity was assessed using the Agilent Bioanalyzer RNA Pico kit. RNA integrity numbers (RIN) ranged from 2.5 to 7.9.

### Visium Tissue Optimization protocol and spatial gene expression

Spatial transcriptomics libraries were generated using the 10x Genomics Visium platform following the manufacturer’s protocols (Visium Tissue Optimization, Rev E; Visium Spatial Gene Expression, Rev F). Optimal permeabilization time was empirically determined for each tissue and standardized to 5 min across samples. cDNA libraries were amplified (13–15 PCR cycles), quality-controlled using Bioanalyzer and Qubit assays, and sequenced on an Illumina NextSeq 2000 platform.

### Spatial V(D)J capture library preparation and long-read sequencing

To enable spatially resolved TCR and BCR profiling, amplified Visium cDNA was subjected to hybridization capture targeting immunoglobulin and T cell receptor sequences using custom xGen Lockdown probes (Integrated DNA Technologies). The protocol largely followed Engblom et al. 2023, with minor adaptions and additional capture probes targeting γδ TCRs. Captured libraries were sequenced on a PacBio Sequel II system, detailed protocols are available upon request.

### Data Analysis of single-cell dataset scRNA-seq preprocessing and integration

Raw gene-cell matrices from seven SNC tumors were processed using Seurat. Cells with fewer than 200 detected genes or more than 10% mitochondrial reads were excluded. To minimize sample-specific feature effects during integration, each dataset was first restricted to the intersecting set of expressed genes (14,003 shared features across tumors) before merging. Counts were log-normalized and mitochondrial percentages regressed out during scaling with *ScaleData*. Highly variable genes were identified using *FindVariableFeatures*, and principal component analysis guided selection of 13 PCs for graph-based clustering and UMAP embedding. Importantly, all TCR and immunoglobulin constant and variable region genes were excluded during dimensional reduction to prevent clonotype identity from driving transcriptional clustering.

### T cell subsetting and reclustering

Following global clustering, CD3E⁺CD3D⁺ T cells were subset and re-embedded using PCA and UMAP. Within this global T cell UMAP, a proliferating cluster (MKI67⁺STMN1⁺TYMS⁺) formed a mixed cycling population containing both CD4⁺ and CD8⁺ cells, consistent with the dominant effect of cell-cycle programs on clustering. For lineage-focused analyses, two independent objects were generated: a CD8⁺ T cell UMAP(CD3E⁺CD3D⁺CD8A⁺CD8B⁺) and a CD4⁺ T cell UMAP (CD3E⁺CD3D⁺CD4⁺).

From the global T cell UMAP, the small “dividing T cells” cluster was included into both lineage-specific objects. Each subset was then reclustered at higher resolution to increase granularity, and minor clusters with discordant lineage markers (for example CD8A⁺ cells within the CD4⁺ object, or vice versa) were iteratively removed, followed by re-embedding until CD8 and CD4 UMAPs were largely free of cross-lineage contamination.

Because the mixed proliferative cluster was intentionally included in both lineage-specific sets, a small number of cells were represented in both CD8⁺ and CD4⁺ objects. Quantitative overlap analysis confirmed that only 59 cells (∼0.87% of CD8⁺ and ∼0.61% of CD4⁺ cells) were shared between objects and were confined to proliferative compartments. Given this low overlap and its confinement to cycling cells, this redundancy is unlikely to affect phenotypic or trajectory conclusions and is consistent with standard practice in single-cell T cell analyses where mixed proliferative clusters are resolved by lineage-specific reclustering.

Finally, full RNA matrices containing TCR variable and constant transcripts, as well as non-shared tumor features, were reintegrated into the lineage-specific CD8 and CD4 objects, re-normalized and re-scaled before downstream V(D)J analyses, ensuring clonotype inference without influencing clustering.

### Sensitivity analysis for sample SNC6

The sample SNC6 was subsequently identified as a cutaneous squamous cell carcinoma with sinonasal involvement. To evaluate the impact of this case on our single-cell results, we repeated the full processing pipeline from raw filtered matrices excluding SNC6, using identical preprocessing parameters and re-annotated clusters to match the original naming. Cluster concordance between the full and SNC6-excluded analyses was assessed using the Adjusted Rand Index (ARI) on cells present in both datasets, a confusion matrix of cluster assignments, and Jaccard overlap of top marker genes (top 50) per cluster.

### Pseudotime and trajectory inference analysis

To reconstruct differentiation trajectories among intratumoral T cells, we performed pseudotime analysis using the Monocle3 package (v1.3.1) in R. T cells were first subsetted from the full scRNA-seq dataset and re-embedded using uniform manifold approximation and projection (UMAP) within Monocle3 to preserve lineage structure. We used the learn_graph function to infer the principal graph structure. The root of the trajectory was manually defined by selecting cells within the Stem-like 1 cluster, based on the expression of naïve/stem-associated markers. Pseudotime values were computed using the order_cells function, and cells were ordered along the trajectory according to their transcriptional similarity and inferred developmental distance from the root. Gene expression changes as a function of pseudotime were identified using the graph_test and fit_models functions.

### Clonal overlap analysis

Clonal overlap between CD8⁺ T cell clusters was quantified using TCR sequence identity derived from paired αβ V(D)J sequencing. For each pair of clusters, overlap was calculated using the overlap coefficient, defined as the number of shared clonotypes normalized by the size of the smaller cluster. Overlap matrices were visualized using a consistent color scale across comparisons to facilitate interpretation of relative clonal sharing between subsets.

### PhyloTrajectory analysis

Count matrices of CD4⁺ and CD8⁺ clonotypes across cell clusters were constructed per patient and subsequently concatenated. Proliferative clusters were excluded from both CD4⁺ and CD8⁺ analyses due to their mixed phenotypic composition. The CD8⁺ TRM/Exh cluster was additionally excluded due to insufficient cell numbers (<50 cells). PhyloTrajectory inference was performed using an Ornstein–Uhlenbeck (OU) process as the proposal distribution for the equilibrium mean and the proposal distribution for the root state; all remaining parameters were set to their default values. Consensus differentiation trees were derived using HIPSTR applied to the MCMC-sampled tree posterior; the full distribution of sampled trees is displayed as background to reflect topological uncertainty. Full details of this method can be found at: https://github.com/MurrellGroup/Phylotrajectories.jl

### Gini coefficients and Shannon diversity

To ensure robust estimation of clonal inequality, we restricted the analysis to cluster-sample combinations containing at least 50 cells to prioritize robustness, reduce noise in rarefaction-based Gini coefficient calculations, and improve the reliability of comparisons across clusters. Lower thresholds increased the number of available cluster-sample combinations but did not materially alter overall trends.

Shannon diversity was calculated to quantify TCR repertoire diversity. Unlike the Gini coefficient, Shannon estimates were less sensitive to rarefaction thresholds and were therefore computed using the same cluster-sample combinations without additional filtering beyond standard quality control.

### Data Analysis of spatial dataset Data preprocessing

Raw fasta files were processed using the 10x Genomics’ Spaceranger count pipeline (version 2.0.1), which performs demultiplexing, barcode processing, alignment to the reference human genome GRCh38-2020-A, and generation of gene expression matrices. When manual alignment was needed it was performed with Loupe 6.0.

Downstream analysis of the spatial gene expression data was performed in R using the semla package and custom scripts. Filtered feature barcode matrices, tissue position files, scale factors and high-resolution H&E tissue image generated from Spaceranger were loaded into a Seurat object for subsequent analysis.

### H&E Image and Gene expression matrix preparation for clustering

Images were masked to remove tissue background and fiducials and then transformed for reorienting the replicate sections to be aligned with each other when plotting. Low quality spots of less than 200 counts or less than 20 genes were filtered out. Hemoglobin genes c("^HBD", "HBG1", "HBG2", "HBE1", "HBZ", "HBM", "HBA2", "HBA1", "HBQ1") were removed from the dataset to avoid blood contamination in the tumor excision borders driving clustering. To control for the effect of certain expanded clones in the clustering, all IG and TCR variable genes in the gene expression matrix were collapsed into 7 major features: IGHVDJ, IGKVJ, IGLVJ, TRAVJ, TRBVDJ, TRGVJ, TRDVDJ.

### Normalization, scaling, PCA dimensionality reduction and clustering

We used the Seurat R package functions for the next steps. Sections were split and log-normalized individually (*NormalizeData*), *FindVariableFeatures* was applied, and then sections were merged back together (union of variable features). The merged dataset was scaled (*ScaleData*), and PCA (*RunPCA*) was performed. The number of PCs that explained at least 90% of cumulative variance was fed into clustering. Cluster resolution was set taking into account what was known about the histopathology and to approximate, although not necessarily match the number of clusters that were automatically generated from the spaceranger count pipeline. For SNC2, res = 0.5 and PCs = 35. For SNC3, res = 0.5 and PCs = 33. For SNC4, res = 0.4 and PCs = 27. Cluster markers were filtered by Avg_log2FC >1, P_val_adj < 0.05.

The top 20 markers per cluster were selected and then tau scores were computed per gene, then genes with tau >= 0.80 were kept, and the top genes were used for generating Supplementary Fig. 1E DotPlots.

Loupe browser files were opened, tumor spots, stroma or normal epithelium (in SNC3) were manually subsetted. Next, DE was run on Loupe browser, and top DE genes from these regions (Fibroblast/Stroma, Tumor, Normal Epithelium) and standard genes expressed by adaptive lymphocytes (*CD3E, CD3D, CD79A, CD79B, JCHAIN*) were plotted in the Fig. 1E DotPlots.

UMAPs were generated using *RunUMAP* from Seurat and feeding the same number of PCs as was determined for clustering.

Integration with Harmony (*RunHarmony*) was tested for all patients, converging in less or equal to 3 iterations and yielding identical clustering results, so it was not used.

### Patient-weighted NMF deconvolution and correlation of factors with clonal diversity

Because NMF minimizes a global reconstruction error, samples contributing larger numbers of spatial spots would otherwise disproportionately influence the factorization. To control for unequal spot numbers and replicate imbalance across patients, we implemented a patient-weighted NMF strategy in which each patient contributed equally to the objective function. Specifically, the expression matrix was split by patient, and spot-level expression values were scaled by a patient-specific weight proportional to 1/ sqrt(nspots) prior to concatenation and NMF. This weighting equalizes each patient’s contribution to the mean-squared reconstruction error while preserving within-patient structure. Non-negative matrix factorization (NMF) was applied (*RunNMF*) to identify latent transcriptional programs with k = 18.

To relate transcriptional programs to local immune repertoire diversity, we quantified clonal richness per spot from the spatial VDJ object and integrated these metrics into the NMF object metadata. For each spot, TRB and IGH richness were defined as the number of distinct non-zero clones (i.e., number of features with counts > 0) computed from the TRB and IGH count matrices. These per-spot richness values were added to the metadata of the NMF Seurat object after matching barcodes between objects.

NMF factor activity per spot was taken from the NMF reduction embeddings (spot loadings). We then computed Spearman rank correlations across spots between each NMF factor loading and each repertoire richness metric (TRB and IGH). Correlations were tested using cor.test(), with settings method="spearman", exact=FALSE, on spots with finite values, and multiple testing across factors was controlled using Benjamini–Hochberg (BH) adjustment. Results were visualized as ranked lollipop plots showing correlation direction and magnitude (ρ) per factor (Fig. 7A).

### Processing of spatial V(D)J long-read data

PacBio HiFi reads in BAM format were demultiplexed by sample index using lima with the symmetric HiFi-preset and an index plate FASTA file, generating one BAM file for each originating Visium library (tissue section). These files were then processed individually using the PacBio Iso-Seq3 pipeline. PCR handles were removed and each read was tagged with a unique molecular identifier (UMI) and spatial barcode. Poly(A) tails were trimmed, and only full-length, non-concatemer reads were retained, by requiring the presence of a poly(A) tail. Spatial barcode correction was performed using a predefined whitelist of Visium barcodes, allowing a maximum edit distance of one. Finally, PCR duplicates were collapsed to generate a FASTA file for each library, which was used as input for V(D)J mapping.

Trimmed, tagged and deduplicated long-read sequences were subsequently annotated using IgDiscover with the run_spatial module. All libraries (n=8, from 3 patients) were processed jointly but separately for TCR and Immunoglobulin annotation. Library identifiers were retained throughout the analysis and spatial barcode information was preserved to enable mapping of clonotypes back to their spatial location (Visium spots). A 250-nt V gene coverage threshold was used for IG and initial TCR annotation. TCR annotation was also repeated with a less stringent 20-nt V gene coverage threshold, which showed high concordance with the 250-nt dataset while increasing recovered UMI counts. The 20-nt dataset was used for visualizing the top expanded clones on tissue, after matching to the scRNA-seq clonotype data.

IgDiscover-derived clonal count matrices were imported in R and added as separate assays to the Seurat object containing histology images and gene expression data.

### Data Analysis of Spatial V(D)J dataset TRB-IGH co-localization analysis

Co-localization analysis was performed separately in tumor and stromal compartments to avoid compartment-driven bias. Spatial spots were classified based on merged spatial gene expression clusters. Within each compartment, spots were scored as TRB-positive or IGH-positive, and co-localization was assessed using the Jaccard index, defined as the number of spots positive for both TRB and IGH divided by the number positive for either. To assess whether the observed overlap was greater than expected by chance, permutation tests were performed by randomly shuffling TRB-positive spots within each compartment 1,000 times. Observed Jaccard indices were then compared with the permutation-derived null distribution.

### Clonotype sharing across spatial clusters

To visualize clonotype sharing across spatial clusters, clonotype presence was defined by binary detection across spatial spots. For each cluster, all clonotypes detected in at least one spot assigned to that cluster were retained. Pairwise sharing was then quantified as the number of clonotypes present in both clusters. Shared clonotype counts were visualized in chord diagrams using the R packages ggplot2 and circlize. For clarity, chord diagrams were focused on sharing between lymphocyte clusters and tumor clusters; other sharings were not shown in the plots.

### Clonal abundance comparison between scRNA-seq and spatial transcriptomics

Clonotype matching between scRNA-seq and spatial transcriptomics was performed based on TCRβ sequences, as TCRβ was consistently and robustly detected in the spatial data. In the scRNA-seq dataset, clonotypes were initially defined using full V(D)J information, including paired TCRα and TCRβ sequences when available. In rare cases, individual scRNA-seq clonotypes exhibited multiple TCRβ transcripts or alternative TCRα pairings; however, these events were infrequent and primarily corresponded to low-frequency clonotypes.

TCRβ clonotypes were independently identified in the scRNA-seq and spatial transcriptomics datasets. For each dataset, clonal abundance was quantified as relative clonal frequency. In the scRNA-seq data, clonal frequency was defined as the proportion of T cells assigned to each clonotype relative to the total number of T cells with productive V(D)J information. In the spatial transcriptomics data, clonal frequency was defined as the proportion of TCRβ-associated transcript counts for a given clonotype relative to the total TCRβ counts across all spatial spots. Clonal frequencies from the two modalities were merged based on shared TCRβ clonotype identifiers, while clonotypes detected in only one modality were retained as modality-specific. To enable visualization on a logarithmic scale, a pseudocount of 1×10⁻⁴, smaller than the minimum observed clonal frequency, was added prior to log₁₀ transformation. Scatter plots were generated comparing clonal frequencies between scRNA-seq and spatial transcriptomics. When multiple clonotypes shared identical frequency values, points were collapsed and the number of clonotypes represented was encoded by point size.

### Processing of blood and tumor tissue for flow cytometry

Healthy or patient human peripheral blood mononuclear cells (PBMCs) were obtained from Karolinska Universitetssjukhuset and isolated by density gradient centrifugation using Lymphoprep following manufacturer’s instructions (Stemcell Technologies). Tumor tissue was collected in PBS and then processed within 1-2 hours after surgery and then processed as described above. In the case of using cryopreserved PBMCs or tumor single cells (TSCs), cells were thawed at 37°C and quickly washed with 37°C cRPMI for a total of two times before staining for flow cytometry. Ethical permission to study PBMCs from healthy donors was obtained from the regional ethical committee in Stockholm (2020-02604).

Immediately after isolation or thawing of PBMCs or TSCs, the cells were transferred to a 96-well v-bottom plate and incubated with Fc Block (BD Biosciences) and LiveDead Fixable Violet (Invitrogen) in PBS for 15 minutes at room temperature in the dark. Cells were washed once with PBS containing 2% FBS and 2.5mM EDTA (FACS buffer) then stained for 30 minutes at room temperature in the dark with the antibody master mix (Contained in supplemental file) containing Brilliant Stain Buffer (BD Biosciences). Cells were washed twice with FACS buffer and then fixed for 30 minutes at room temperature with eBioscience Foxp3 / Transcription factor staining kit (Invitrogen). Following fixation, cells were washed twice with appropriate permeabilization buffer and then incubated overnight (18 hours) with intracellular antibody master mix containing Fc Block (BD Biosciences) at 4°C. Cells were washed with FACS buffer and then resuspended in PBS and kept at 4°C for no more than 1 day until acquisition. Single-color controls were done using cells when possible or UltraComp eBeads Plus Compensation Beads (Invitrogen) with the same methodology as described above. All samples were acquired on a 5-laser ID7000 Spectral Cell Analyzer (Sony) following daily passing of the quality control using Align Check beads (Sony). Unmixing, when needed, was done manually using ID7000 Analysis Software (Sony). Analysis and gating of flow cytometry data was done using Flowjo v10.7 (BD Biosciences). Flowjo exported csv files containing frequencies of populations were plotted using python v3.10.13, pandas v2.2.3, and seaborn v0.13.2 in the case of heatmaps or GraphPad Prism v10.6.1 (Dotmatics) in the case of bar graphs.

## Competing interests

K.T. and C.E. are inventors on the patent (US Patent no. US11692218B2) encompassing the spatial VDJ work.

## Data availability

The single-cell RNA sequencing (scRNA-seq) and V(D)J sequencing data generated in this study have been deposited in the Swedish National Data Service (SND) repository. The data include raw gene expression matrices and associated metadata at both the cell and sample levels. Access to the dataset will be provided upon reasonable request, in accordance with ethical and data protection regulations.

## Acknowledgements

The authors would like to acknowledge numerous funding bodies, including Vetenskapsrådet, Cancerfonden, LEO Foundation, NovoNordisk Foundation and Carlsberg Foundation. Thank you to Nicholas Huntington for critical review of the manuscript.

## Supplementary File

### Supplementary Figures

**Supplementary Figure 1.**
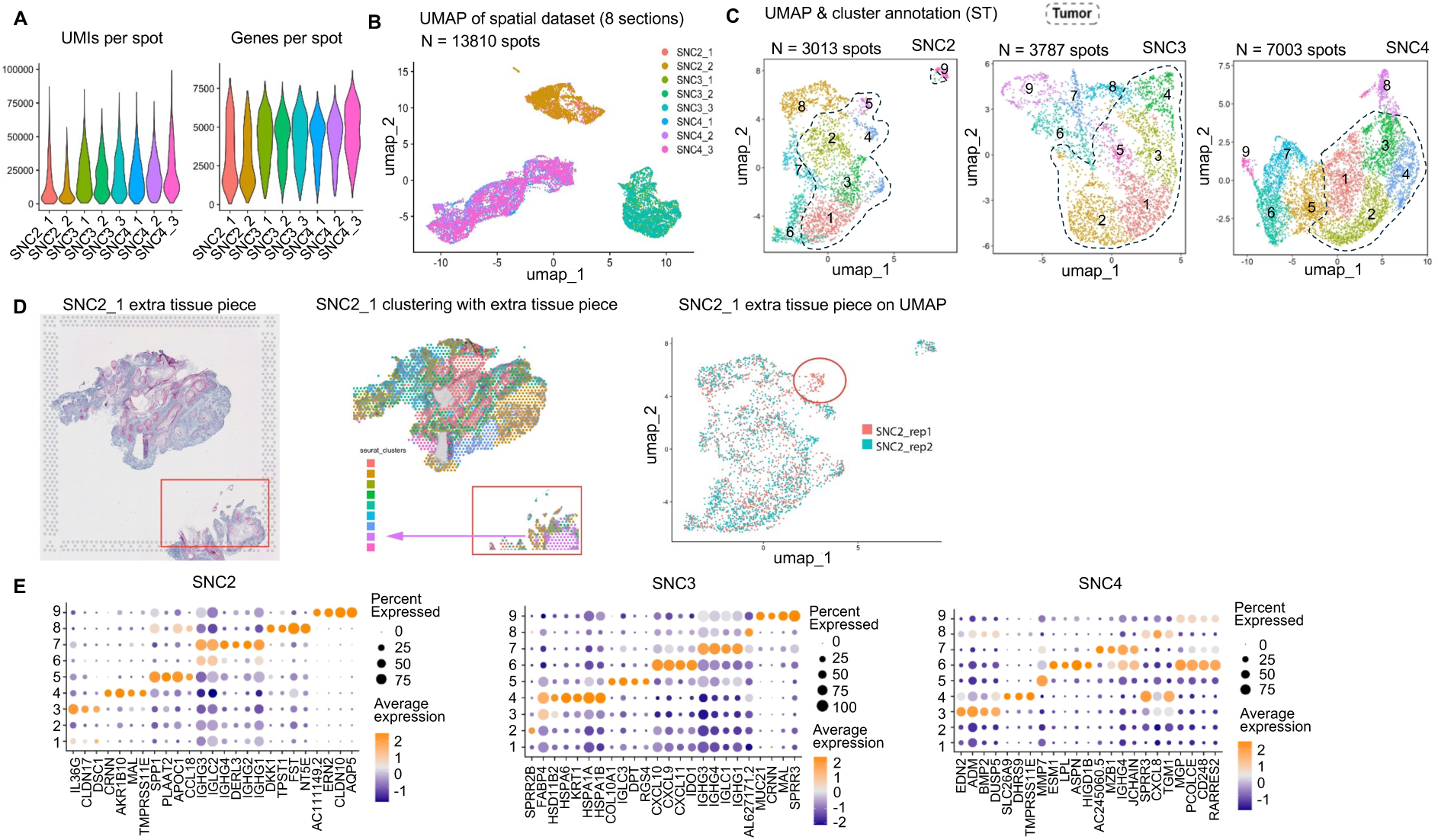
Spatial transcriptomics reveals diverse immune and non-immune compartments in SNC. (**A**) Violin plots showing the distribution of UMIs and genes detected per Visium spot across all tumor sections. (**B**) UMAP representation of the spatial transcriptomics dataset (13,810 spots from 8 sections), colored by section. (**C**) UMAPs of SNC 2-4 spatial transcriptomic data colored according to spatial cluster identities. The dashed border groups clusters most representative of tumor epithelium. (**D**) Display of an additional tissue fragment from patient SNC2. Left: H&E image of SNC2_rep1 with the extra fragment indicated by a box; middle: clustering of SNC2_rep1 including the extra fragment, showing that the purple cluster predominantly maps to this region; right: UMAP of all SNC2 spots, with spots corresponding to the extra tissue fragment highlighted in a red circle. (**E**) Dot plots displaying the average expression and percent of spots expressing up to six top markers per cluster with high specificity (tau ≥ 0.8), minimum detection in 25% of spots, minimum log2 fold change >1, and p adjusted value ≥ 0.01.

**Supplementary Figure 2.**
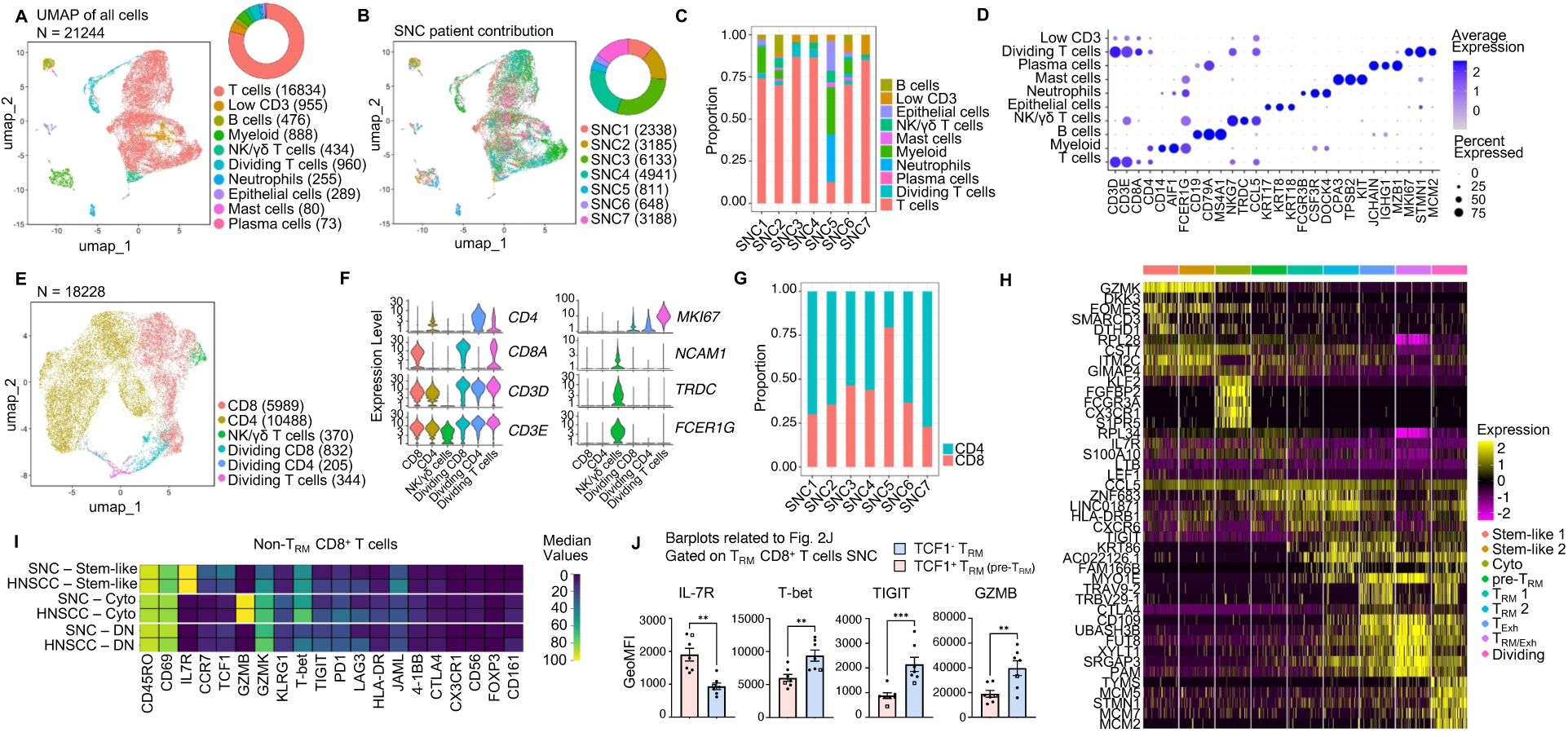
Single-cell RNA-Seq of SNC identifies CD8⁺ T cells of diverse transcriptional states. (**A**) UMAP visualization of integrated scRNA-seq dataset from seven SNC tumors (n = 21,244 cells). Donut chart of the relative frequencies of identified clusters. (**B**) UMAP colored by patient. Donut chart indicates the proportion of cells from each tumor. (**C**) Barplot of cell-type composition per patient in the single-cell dataset. All SNC samples included diverse immune and non-immune populations, though relative abundance varied across tumors. Most tumors, apart from SNC5 and SNC6 were biased to profile T cells, although non-T cells were always included in the purification for all tumors. (**D**) DotPlot depicting the average expression and percent of cells expressing canonical markers across major cell types in the single-cell dataset. (**E**) UMAP visualization of 18,228 CD3⁺ T cells from SNC tumors. (**F**) Violin plots of representative markers expression across CD3⁺ T cells clusters. (**G**) Proportion of CD4⁺ and CD8⁺ T cells per patient. (**H)** Heatmap of top 5 genes expressed across CD8⁺ T cell clusters illustrating gradual and overlapping transcriptional transitions among TRM and exhausted populations, rather than sharply segregated states. (**I**) Heatmap of markers measured by flow cytometry in tumor-infiltrating CD8⁺ T cell subsets. Cells were gated from non-TRM CD8⁺ (CD69⁺ CD103⁺) and classified as stem-like (GZMB⁺IL7R⁺), cytotoxic (GZMB⁺IL7R⁺) or double negative (DN; GZMB⁺IL7R⁺) cells. For each subset, SNC and HNSCC samples are shown separately. Values represent the median percentage across SNSCC (n = 7) or HNSCC (n = 8) tumors. (**J**) Flow cytometric analysis of TRM cells in SNC tumors, identifying different marker expression in TCF1⁺ and TCF1⁻ TRM populations. Bar plots show the geometric mean fluorescence intensity (MFI) of selected markers (see Fig. 2J). Bars represent mean ± SEM, and individual points correspond to biological replicates; the square indicates SNC6, a cutaneous carcinoma with nasal extension. Statistical significance was assessed using a paired t-test.

**Supplementary Figure 3.**
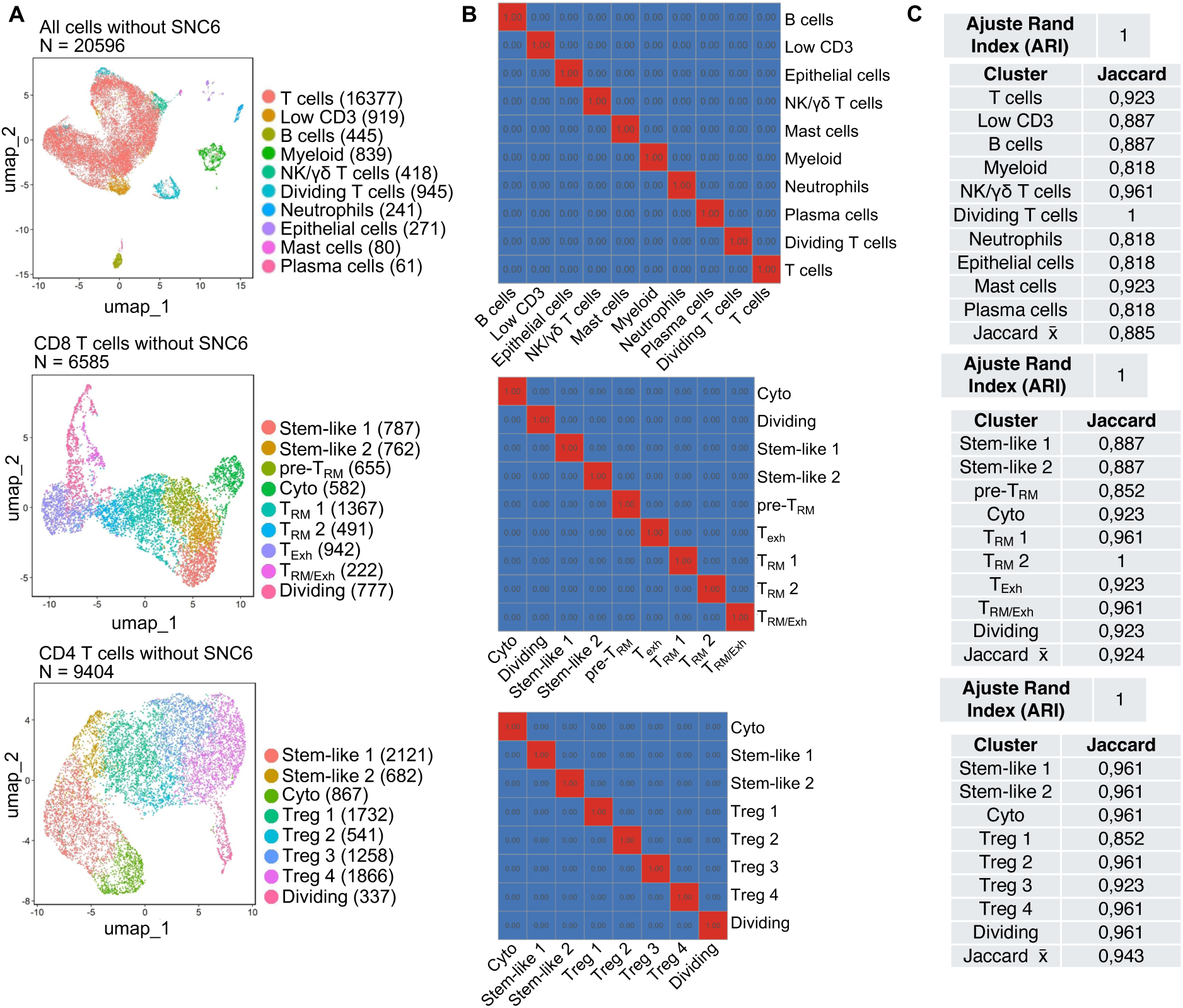
Sensitivity analysis: impact of excluding SNC6 from the dataset is negligible. (**A**) UMAP for all cells (top), CD8 T cells (middle) and CD4 T cells (bottom) without sample SNC6. (**B**) Confusion matrix (normalized by original-cluster size) showing correspondence between cluster assignments in the full analysis (7 samples) and the re-clustered dataset excluding SNC6 (6 samples). Values indicate the proportion of cells from each original cluster that fall into each new cluster. (**C**) Adjusted Rand Index (ARI) quantifying global cluster agreement (ARI = 1.0), and table: per-cluster Jaccard overlap of top 50 marker genes (mean shown), indicating preservation of molecular signatures across the two analyses.

**Supplementary Figure 4.**
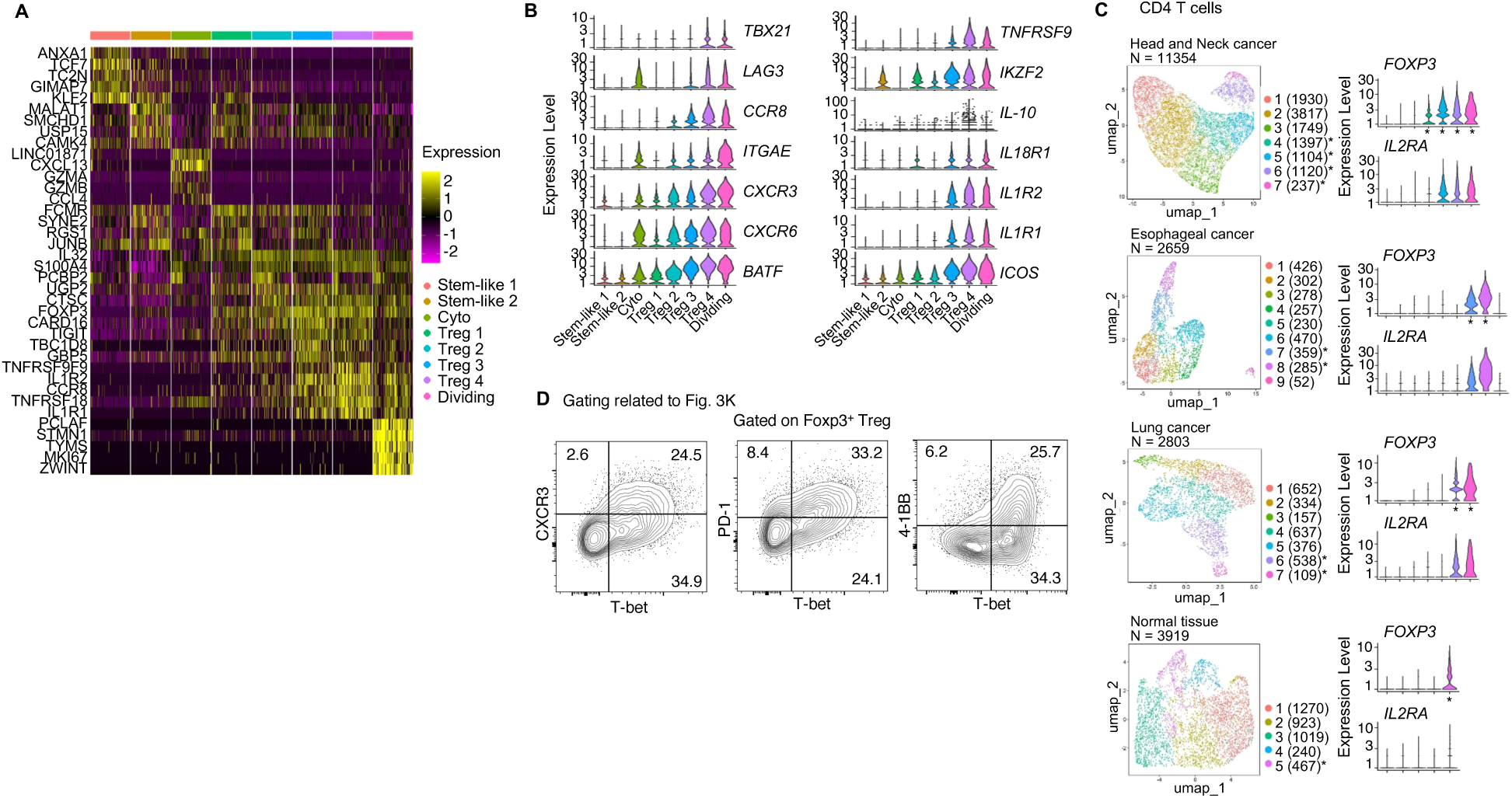
CD4⁺ T cells in SNC tumors exhibit a predominantly regulatory phenotype which shows signs of tumor adaptation. (**A**) Heatmap of top 5 genes expressed across CD4⁺ T cell illustrates gradual and overlapping transcriptional transitions among Treg cell clusters rather than sharply segregated states. (**B**) Violin plots show expression of selected markers across CD4⁺ clusters. Many markers are gradually increased from Treg 1 to Treg 4 state. (**C**) UMAP projections of CD4⁺ T cells from head and neck squamous cell carcinoma (HNSCC; GSE173468), esophageal carcinoma (GSE196756), non-small cell lung cancer (GSE198099), and adjacent normal tissue (GSE173468), following uniform reprocessing of raw scRNA-seq data using the same analytical pipeline applied to SNC. Violin plots show the expression of *FOXP3* and *IL2RA* across clusters in each dataset. Clusters were classified as Treg populations based on detectable FOXP3 expression, independent of original study annotations, and are indicated by asterisks in the UMAPs and corresponding violin plots. This standardized approach enabled direct comparison of Treg frequencies across datasets (see Fig. 3F). (**D**) Representative contour plots of T-bet vs. CXCR3, PD-1 and 4-1BB in gated FOXP3⁺ Treg cells. Linked to bar plots in Fig. 3K.

**Supplementary Figure 5.**
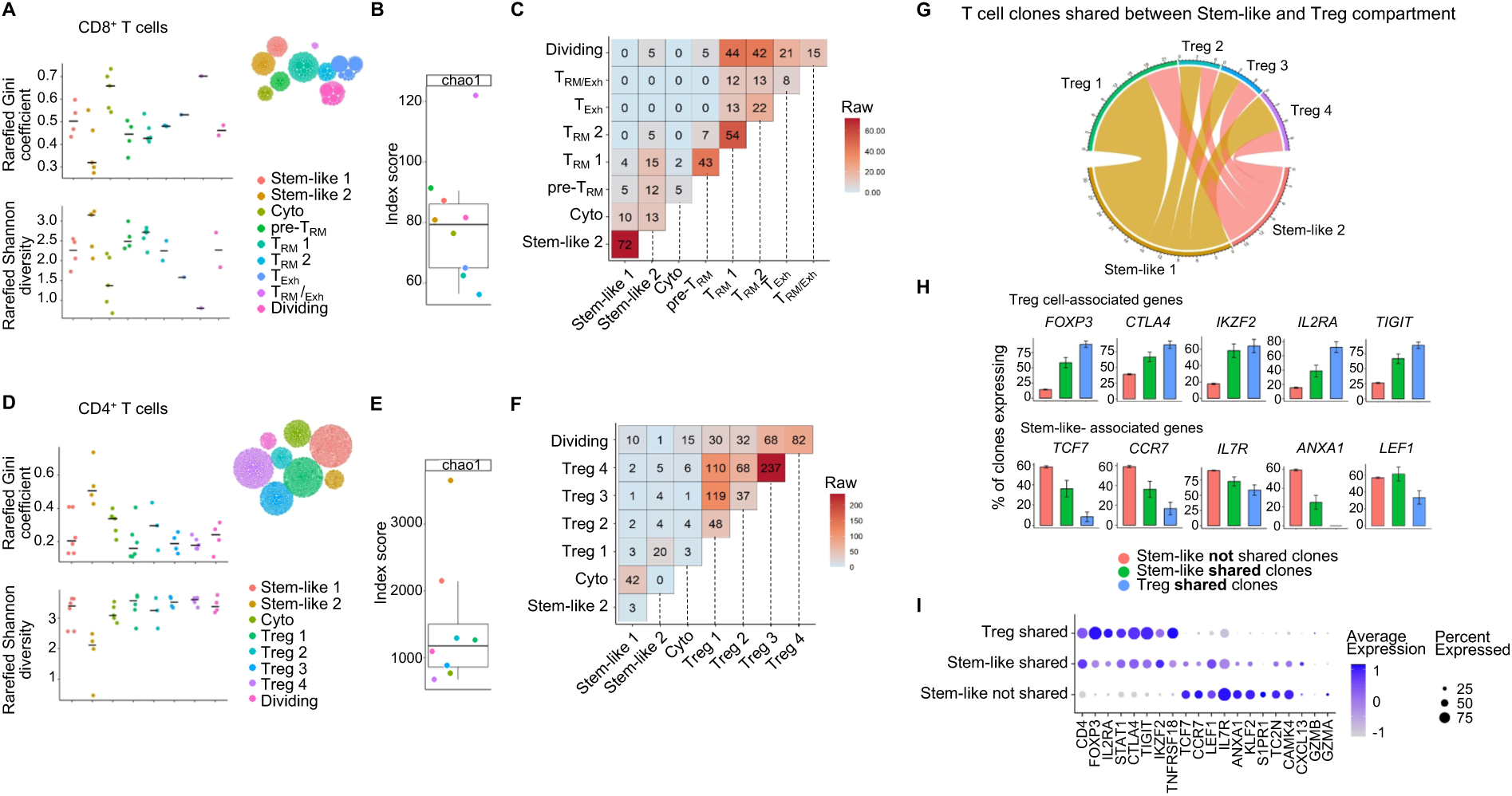
Clonal profiles of CD4^+^ and CD8^+^ T cells in SNC. (**A**, **D**) Rarefied Gini coefficients (top) and Shannon diversity indices (bottom) across CD8⁺ (A) and CD4⁺ (D) T cell clusters. (**B**, **E**) Chao1 richness estimates for CD8⁺ (B) and CD4⁺ (E) T cell clusters. (**C**, **F**) Pairwise clonal overlap between CD8⁺ (C) and CD4⁺ (F) T cell clusters shown as raw shared clone counts. (**G**) Chord diagram showing shared clonotypes between stem-like (Stem-like 1-2) and Treg (Treg 1-4) clusters within CD4⁺ T cells from single cell data. Chord width reflects the number of shared clonotypes between connected populations. Analysis was restricted to clonotypes present in both stem-like and Treg compartments. For clarity, only inter-compartment interactions are shown, while intra-compartment connections (stem-like–stem-like or Treg–Treg) and interactions involving other clusters were excluded. (**H**) Bar plots showing the proportion of clonotypes expressing selected Treg-associated and stem-like-associated genes across CD4⁺ T cell compartments. Clonotypes were grouped as: *stem-like not shared* (clones restricted to stem-like clusters), *stem-like shared* (clones in the stem-like compartment shared with Treg clusters), and *Treg shared* (clones in the Treg compartment shared with the stem-like clusters). For each gene, bars represent the percentage of clonotypes in which the majority (>50%) of cells express the indicated gene. Error bars denote standard error of the mean across clonotypes. (**I**) Dot plot depicting the average expression and percent of cells expressing selected markers across *stem-like not shared*, *stem-like shared*, and *Treg shared* cells within CD4⁺ T cells.

**Supplementary Figure 6.**
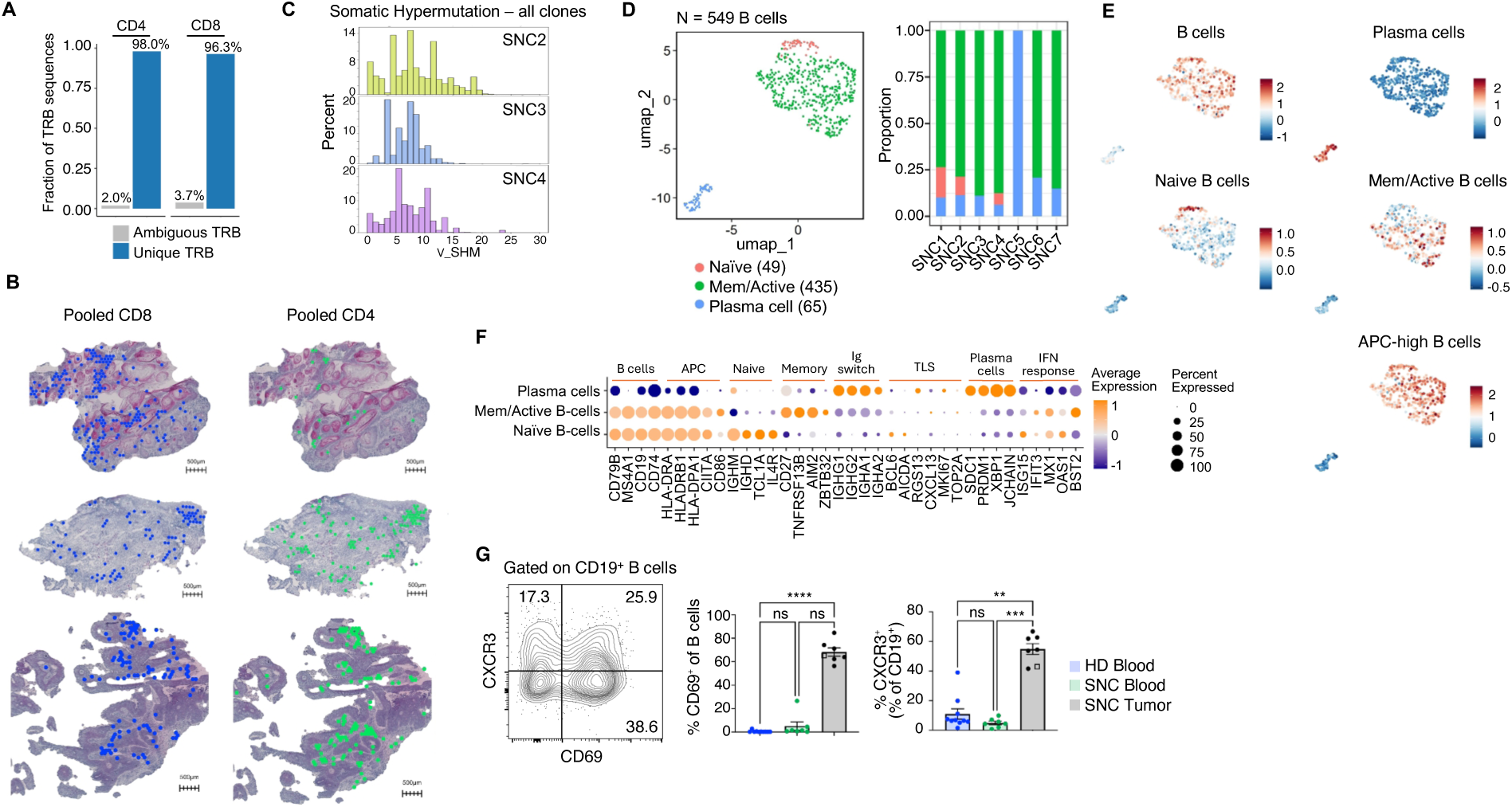
Activated B cells do not show a TLS phenotype in SNC. (**A**) Bar plots show the fraction of unique TCRβ (TRB) CDR3 nucleotide sequences that map to a single clonotype (“Unique”) or to multiple clonotypes (“Ambiguous”), as defined by paired TRA–TRB information in the scRNA-seq dataset. (**B**) Pooled CD8⁺ and CD4⁺ T cell clones plotted on tumor sections. Colored spots indicate at least one UMI count from a clonotype annotated as CD8^+^ or CD4^+^ based on the corresponding single cell data. Scale bars, 500 µm. (**C**) Histograms showing the degree of somatic hypermutation (SHM) in B cell clonotypes for each SNC2-4 tumor. Only clonotypes supported by more than one UMI were included to reduce technical noise. SHM is expressed as the percentage of nucleotide divergence from the inferred germline IGH sequence. (**D**) UMAP of tumor-infiltrating B and plasma cells from SNC patients resolved into three subsets: naive B cells, memory/activated B cells, and plasma cells (left), and distribution of B cell states across individual SNC tumors (right). (**E**) UMAP projections of B-lineage cells showing module scores for curated gene signatures corresponding to B cells, naïve B cells, memory/activated B cells, APC-high B cells, and plasma cells. Colors indicate relative enrichment of each gene signature per cell (scaled expression). (**F**) Dot plot showing the average expression and percent of cells expressing selected markers across B cell subsets. (**G**) Flow cytometric analysis of CXCR3 and CD69 expression in CD19⁺ B cells from healthy donor blood, SNC patient blood, and SNC tumors. Representative plots (left) and quantitative summaries (right) are shown. Bars represent mean ± SEM; statistical significance was assessed using a Kruskal–Wallis test followed by multiple-comparison correction. The square indicates SNC6.

**Supplementary Figure 7.**
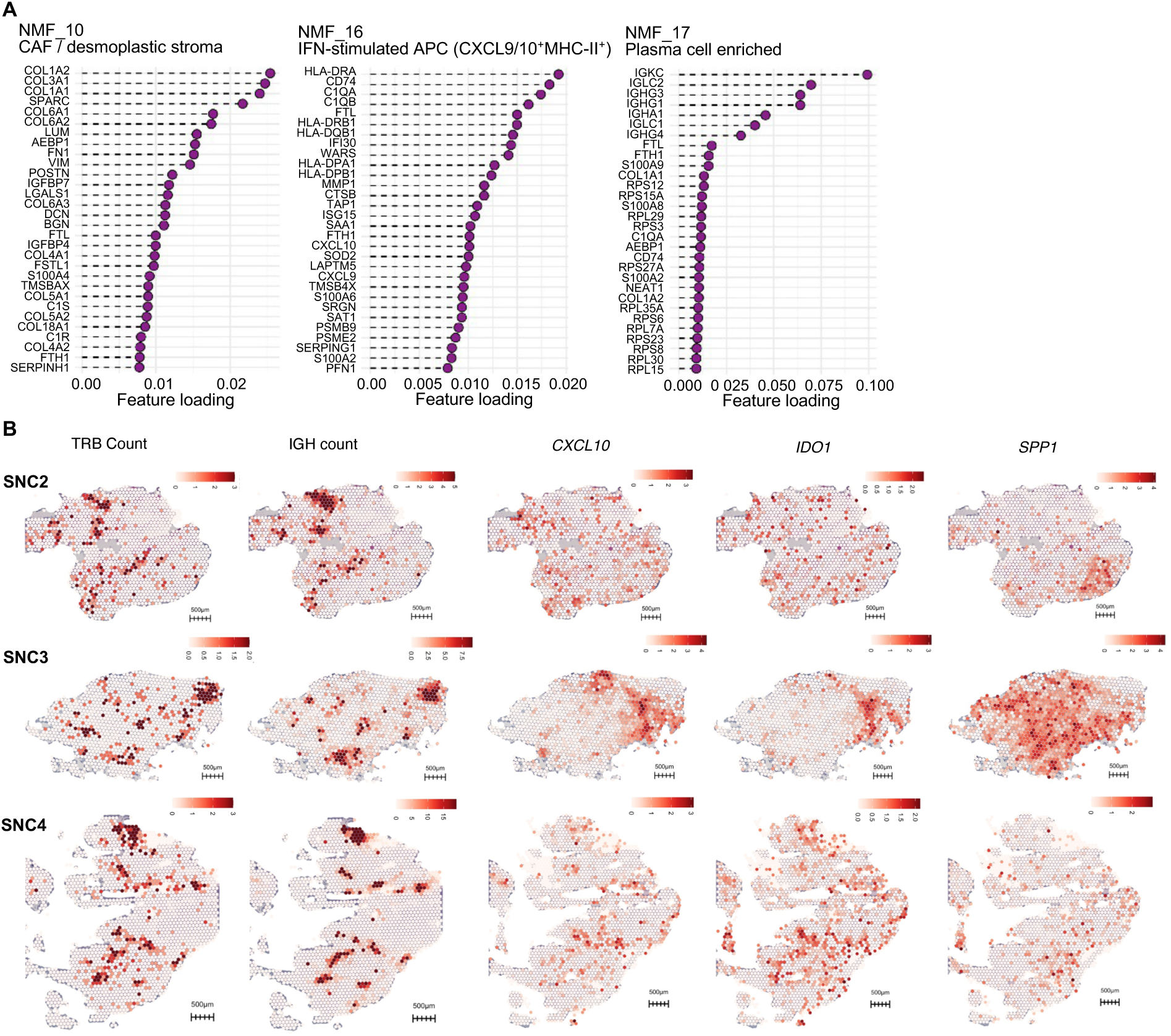
NMF_10, _16 and _17 are associated with T and B cell clones in SNC. (**A**) Top 30 feature loadings for the three diversity-associated NMF programs. (**B**) Spatial maps of total TRB and IGH UMI counts from Spatial V(D)J profiling, and CXCL10, IDO1 and SPP1 expression across tumors. Color scale values indicate UMI counts. Scale bars, 500 µm.

**Supplementary Figure 8.**
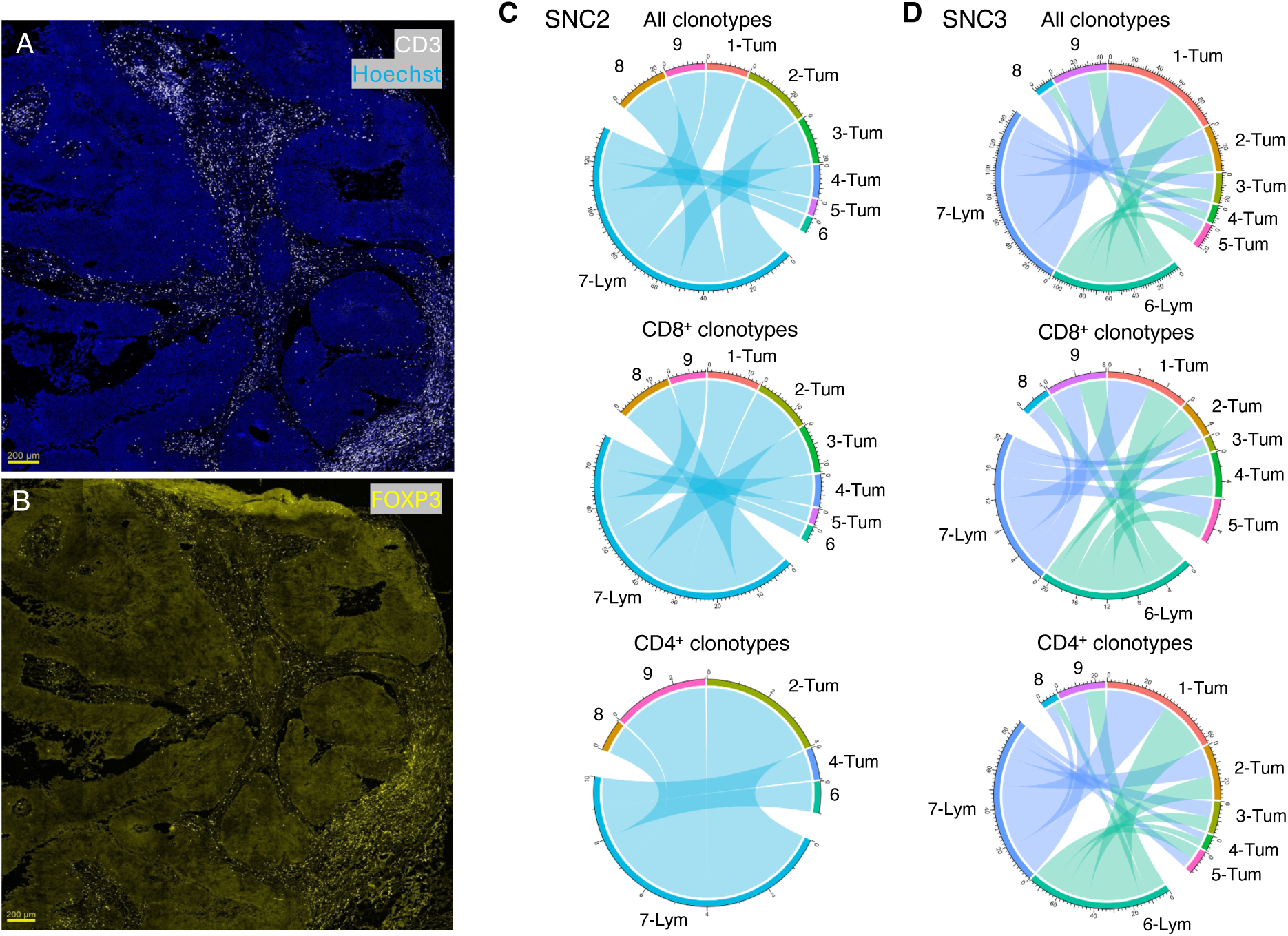
Extensive T cell infiltration in SNC tumors. (**A**) Representative wide-scan confocal image of an SNC tumor section showing spatial distribution of CD3⁺ T cells (white) and nuclei (Hoechst, blue). CD3⁺ T cells are enriched in stromal regions surrounding tumor epithelial areas. (**B**) Corresponding section stained for FOXP3 (yellow), revealing a similar spatial distribution of FOXP3⁺ T cells within the TME. Both panels A and B show low-magnification views of the same tumor region. Scale bars, 200 μm. (**C**, **D**) Chord diagrams of shared clonotypes between lymphocyte-rich clusters (SNC2: cluster 7; SNC3: clusters 6 and 7) and other spatial clusters from spatial transcriptomic data. Shown are ‘all clonotypes’ as aligned by spatial V(D)J, and specific CD8^+^ and CD4^+^ clones, which could be verified by scRNA-seq. For clarity, only inter-compartment interactions are shown, while intra-compartment connections (lymphoid-lymphoid or tumor-tumor) are excluded. These diagrams reveal extensive clonal sharing between lymphoid-rich niches and tumor regions, consistent across both CD8⁺ and CD4⁺ T cell compartments.

### Supplementary Tables

**Supplementary Table 1:**
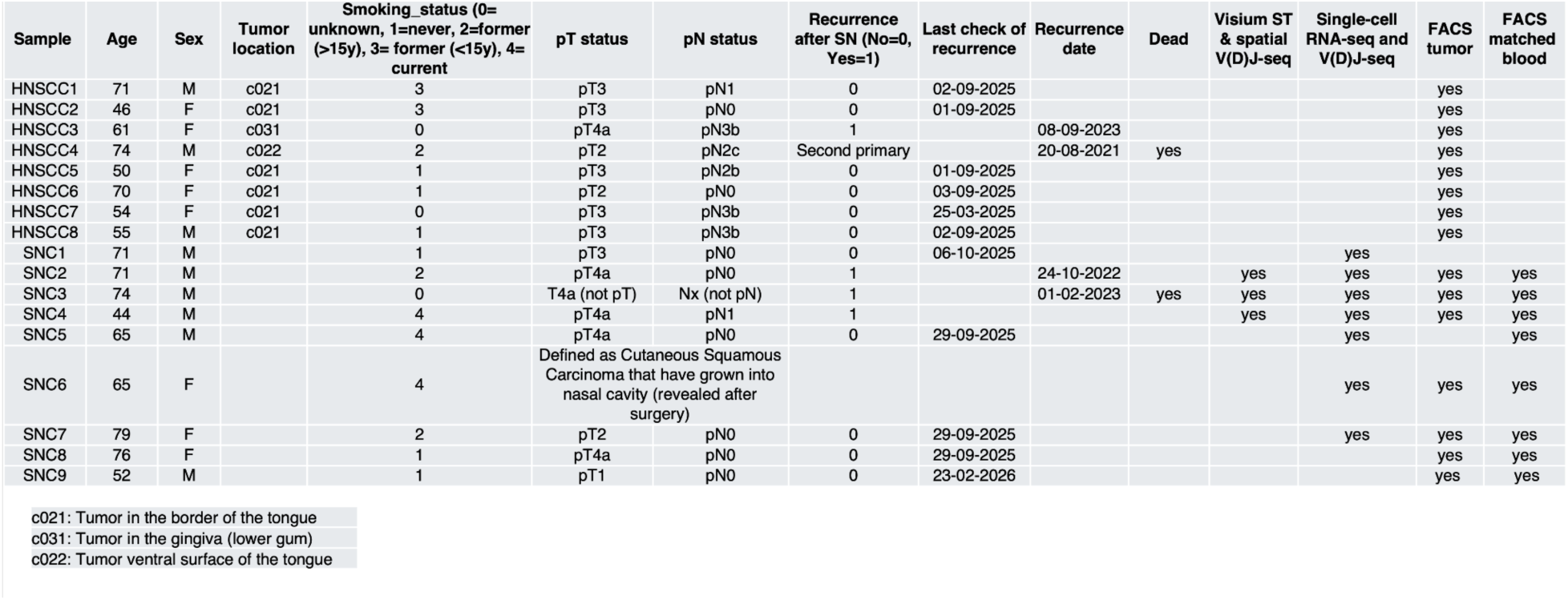
Patient characteristics.

**Supplementary table 2.**
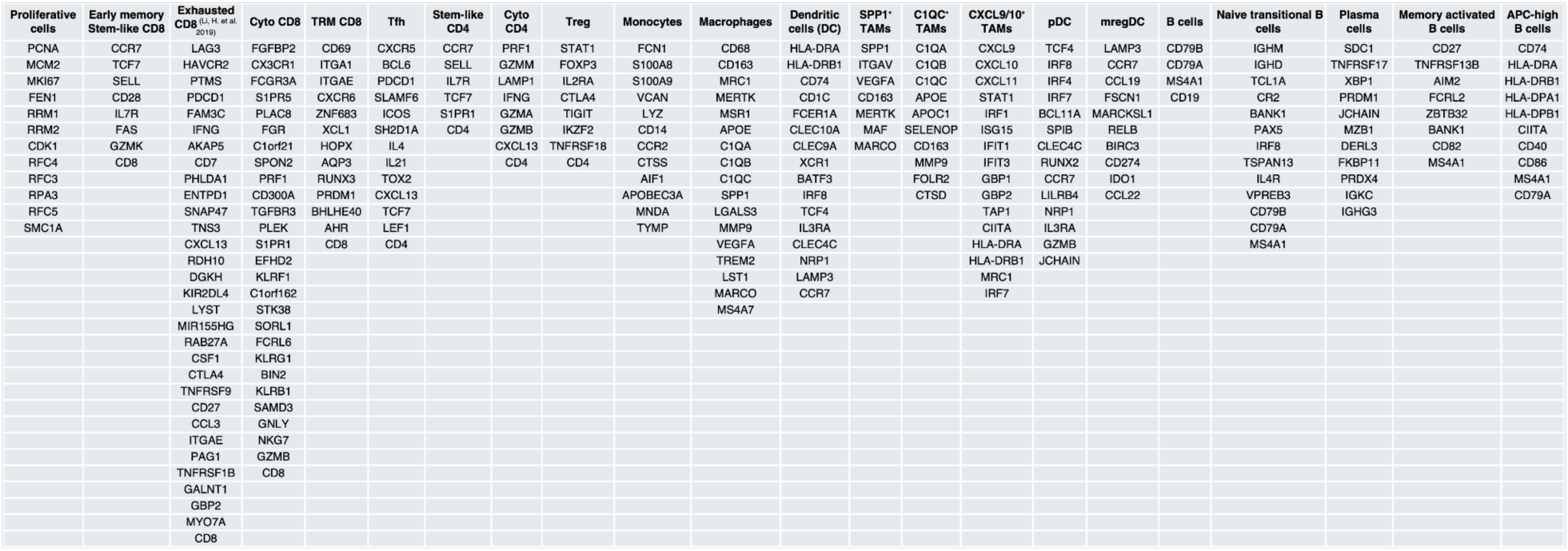
Gene signature used for *AddModuleScore*.

**Supplementary table 3.**
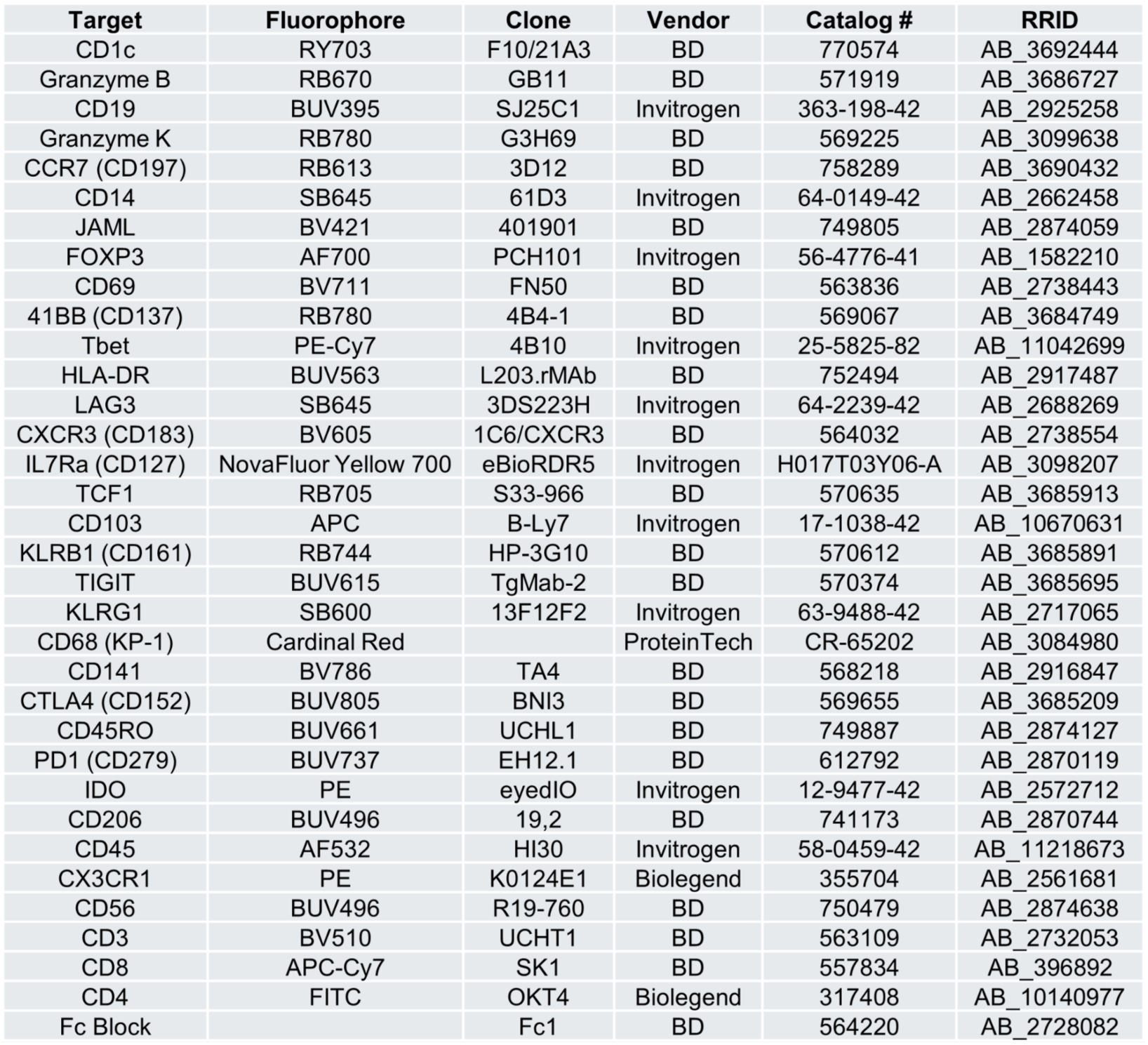
List of antibodies used in the study.

| Target | Fluorophore | Clone | Vendor | Catalog # | RRID |
| --- | --- | --- | --- | --- | --- |
| CD1c | RY703 | F10/21A3 | BD | 770574 | AB_3692444 |
| Granzyme B | RB670 | GB11 | BD | 571919 | AB_3686727 |
| CD19 | BUV395 | SJ25C1 | Invitrogen | 363-198-42 | AB_2925258 |
| Granzyme K | RB780 | G3H69 | BD | 569225 | AB_3099638 |
| CCR7 (CD197) | RB613 | 3D12 | BD | 758289 | AB_3690432 |
| CD14 | SB645 | 61D3 | Invitrogen | 64-0149-42 | AB_2662458 |
| JAML | BV421 | 401901 | BD | 749805 | AB_2874059 |
| FOXP3 | AF700 | PCH101 | Invitrogen | 56-4776-41 | AB_1582210 |
| CD69 | BV711 | FN50 | BD | 563836 | AB_2738443 |
| 41BB (CD137) | RB780 | 4B4-1 | BD | 569067 | AB_3684749 |
| Tbet | PE-Cy7 | 4B10 | Invitrogen | 25-5825-82 | AB_11042699 |
| HLA-DR | BUV563 | L203.rMAb | BD | 752494 | AB_2917487 |
| LAG3 | SB645 | 3DS223H | Invitrogen | 64-2239-42 | AB_2688269 |
| CXCR3 (CD183) | BV605 | 1C6/CXCR3 | BD | 564032 | AB_2738554 |
| IL7Ra (CD127) | NovaFluor Yellow 700 | eBioRDR5 | Invitrogen | H017T03Y06-A | AB_3098207 |
| TCF1 | RB705 | S33-966 | BD | 570635 | AB_3685913 |
| CD103 | APC | B-Ly7 | Invitrogen | 17-1038-42 | AB_10670631 |
| KLRB1 (CD161) | RB744 | HP-3G10 | BD | 570612 | AB_3685891 |
| TIGIT | BUV615 | TgMab-2 | BD | 570374 | AB_3685695 |
| KLRG1 | SB600 | 13F12F2 | Invitrogen | 63-9488-42 | AB_2717065 |
| CD68 (KP-1) | Cardinal Red |  | ProteinTech | CR-65202 | AB_3084980 |
| CD141 | BV786 | TA4 | BD | 568218 | AB_2916847 |
| CTLA4 (CD152) | BUV805 | BNI3 | BD | 569655 | AB_3685209 |
| CD45RO | BUV661 | UCHL1 | BD | 749887 | AB_2874127 |
| PD1 (CD279) | BUV737 | EH12.1 | BD | 612792 | AB_2870119 |
| IDO | PE | eyedIO | Invitrogen | 12-9477-42 | AB_2572712 |
| CD206 | BUV496 | 19,2 | BD | 741173 | AB_2870744 |
| CD45 | AF532 | HI30 | Invitrogen | 58-0459-42 | AB_11218673 |
| CX3CR1 | PE | K0124E1 | Biolegend | 355704 | AB_2561681 |
| CD56 | BUV496 | R19-760 | BD | 750479 | AB_2874638 |
| CD3 | BV510 | UCHT1 | BD | 563109 | AB_2732053 |
| CD8 | APC-Cy7 | SK1 | BD | 557834 | AB_396892 |
| CD4 | FITC | OKT4 | Biolegend | 317408 | AB_10140977 |
| Fc Block |  | Fc1 | BD | 564220 | AB_2728082 |

**Supplementary Note 1. Histopathological and spatial transcriptomic annotation of profiled SNC tumors**

SNC2 was classified as a keratinizing squamous cell carcinoma and contained tumor islands with characteristic keratin pearl formation, indicative of well-differentiated regions, alongside less-differentiated, less-keratinizing tumor islands interspersed with stromal regions. Spatial transcriptomic clustering identified five transcriptionally-distinct clusters associated with keratinizing tumor regions (clusters 1-5), supported by tumor-associated marker expression (Fig. 1E, Supplementary Fig. 1E). Cluster 5 corresponded to a small tissue fragment present only in one replicate section (Supplementary Fig. 1D). Lymphocyte populations including T and B-cells/plasma cells were enriched around keratinizing tumor areas and along stromal corridors, as indicated by expression of *CD3* chains and *CD79A*/*JCHAIN*, respectively. Cancer-associated fibroblast (CAF) signatures, characterized by *POSTN* and *THY1* expression, were primarily detected with lymphoid markers rather than tumor epithelial clusters (Fig. 1E).

SNC3 was a poorly differentiated non-keratinizing advanced-stage cancer. Histological examination identified a small patch of normal epithelium adjacent to areas enriched in lymphoid infiltrates and macrophages, together with widespread tumor infiltration. Consistent with this annotation, spatial clustering resolved a region of normal epithelium expressing *KRT4*, *KRT6C* and *TMEM45B* (cluster 9). In contrast, tumor regions, represented by clusters 1–5, upregulated *MMP13*, *MMP7* and *TREM1*. A lymphoid-rich region located between the normal epithelium and tumor regions expressed genes associated with T and B cells (clusters 6 and 7). Fibroblast-associated genes, including *COL1A1* and *THY1*, were again associated with lymphoid-rich regions, particularly regions expressing B cell-related genes, but were also detected in tumor-associated regions, especially cluster 5 (Fig. 1E).

SNC4 was classified as a poorly differentiated, non-keratinizing tumor and showed a papillary tumor architecture surrounded by stromal areas densely infiltrated by immune cells. Spatial clustering resolved multiple tumor epithelium areas (clusters 1-4), supported by epithelial and tumor-associated marker expression (Fig. 1E, Supplementary Fig. 1E). Several stromal compartments were also identified. Among these, clusters 6 and 7 were strongly associated with T and B cell marker genes, including *CD3* genes, *CD79A* and *JCHAIN*, consistent with lymphocyte-rich stromal regions. Fibroblast/stromal marker expression was mainly detected in non-tumor regions and overlapped with immune-enriched areas, whereas clusters 5 and 9 showed no obvious association with either tumor epithelial or lymphocyte gene programs.

